# Interleukin-33 regulates metabolic reprogramming of the retinal pigment epithelium in response to immune stressors

**DOI:** 10.1101/2021.02.04.429634

**Authors:** Louis M Scott, Emma E Vincent, Natalie Hudson, Chris Neal, Nicholas Jones, Ed Lavelle, Matthew Campbell, Andrew P Halestrap, Andrew D Dick, Sofia Theodoropoulou

## Abstract

It remains unresolved how retinal pigment epithelial (RPE) cell metabolism is regulated following immune activation to maintain retinal homeostasis and retinal function. We exposed RPE to several stress signals, particularly toll-like receptor stimulation, and uncovered an ability of RPE to adapt their metabolic preference on aerobic glycolysis or oxidative glucose metabolism in response to different immune stimuli. We have identified interleukin-33 (IL-33) as a key metabolic checkpoint that antagonises the Warburg effect to ensure the functional stability of the RPE. The identification of IL-33 as a key regulator of mitochondrial metabolism suggests roles for the cytokine that go beyond its extracellular “alarmin” activities. IL-33 exerts control over mitochondrial respiration in RPE by facilitating oxidative pyruvate catabolism. We have also revealed that in the absence of IL-33, mitochondrial function declines and resultant bioenergetic switching is aligned with altered mitochondrial morphology. Our data not only sheds new light in the molecular pathway of activation of mitochondrial respiration in RPE in response to immune stressors, but also uncovers a novel role of nuclear intrinsic IL-33 as a metabolic checkpoint regulator.

## Introductions

The retina is referred to as an “immune privileged” tissue, where immune homeostasis is maintained through hemopoietic immune cells such as microglia and also immune-competent tissue resident cells such as the retinal pigment epithelium (RPE) [1]. The RPE is highly differentiated, and considered to be a post-mitotic single cell layer that performs a host of functions critical to retinal homeostasis, including the maintenance of the visual cycle and photoreceptor phagocytosis [2]. RPE cells are highly metabolically active whilst providing critical metabolic support through, for example, the directional transport of glucose and lactate to fuel the outer retina and photoreceptors in particular [3]. Studies of cultured RPE cells show that they can derive energy from glucose by either aerobic glycolysis or oxidative phosphorylation (OXPHOS), depending upon culture conditions. Defects in RPE metabolic and mitochondrial function, associated with low grade inflammation or with age-related decline, are associated with retinal degenerative diseases including age-related macular degeneration (AMD) [4]. However, the causal link between RPE metabolic alterations and retinal function and disease remains unclear.

Immune cells with different functions use several different metabolic pathways to generate adequate levels of energy stores to support survival and to produce numerous biosynthetic intermediates to allow for cellular growth and proliferation [5]. Understanding how immune-competent tissue-resident cells, like RPE, manage energy consumption to potentially support the maintenance of outer retinal function is crucial, thus unknown. Recently, it has been shown that RPE cells utilise reductive carboxylation [6] to support redox homeostasis. This in turn is re-inforced by the documented disproportionate damage to mitochondrial DNA in the RPE of individuals with AMD. Furthermore, the role of lipids, which accumulate in the macula, and their oxidation, has emerged as an important factor in AMD development [7].

Interleukin-33 (IL-33), a member of the IL-1 family, is a type 2 cytokine that is constitutively expressed in the inner retina, RPE and choroid [8], as well as in epithelial cells, endothelial cells, and fibroblasts [9]. IL-33 functions as an !alarmin” molecule released from barrier cells following cellular damage [10]. IL-33 is a pleotropic cytokine with effects that are either pro- or anti-inflammatory depending on disease or cell context [11]. This is reflected by major roles in infection, allergic responses, metabolic homeostasis and cancer [12–14]. There is growing evidence suggesting that IL-33 plays an important role in systemic metabolic diseases, like type 2 diabetes [15], as well as in cardiac disease [16].

We have previously shown that Toll-like receptors (TLR) ligands (TLR-3,-4) induce up-regulation of IL-33 in RPE cells without cell death. We also demonstrated a protective role for IL-33 in regulating ocular tissue responses to promote wound healing [8]. Recent exploitation of the cytoplasmic hybrid technique for representative mtDNA haplotypes of AMD populations, revealed alterations in IL-33 expression and altered bioenergetics [17]. This, coupled with novel epigenetic roles for IL-33 as shown in the maintenance of mitochondrial function in adipose tissue [18], led us to investigate whether IL-33 acts as an intracellular check point for regulation of RPE metabolism. Here, we characterize RPE metabolic reprogramming in response to TLR ligation. We provide evidence that immune-mediated signals drive differential bioenergetic sourcing within the RPE, leading to a specific nuclear signature of IL-33 expression that maintains mitochondrial health and allows cells to respond and deliver energy intermediates required for RPE function.

## Results

### RPE has a differential metabolic response to varying TLR agonists

The bioenergetic response of RPE cells to potentially harmful signals (through TLR stimulation) was investigated to ascertain the response and ability to maintain function under acute immune stress. We employed a Seahorse XF Analyser to determine whether TLR activation altered RPE cellular metabolism.

Stimulation of RPE by LPS, a potent ligand of TLR4, promoted a significant increase (1.5-fold) in the extracellular acidification rate (ECAR) relative to resting controls (Figure 1A) and this was accompanied by a decrease in oxygen consumption rate (OCR) in human RPE cells (ARPE-19) (Figure 1B). These data imply increased aerobic glycolysis in LPS stimulated ARPE-19 cells (Figure 1C) and this was confirmed by an increase in glucose consumption (Figure 1D). In contrast to LPS, stimulation of RPE by Poly (I:C), a ligand for TLR3, for 6h led to increased aerobic metabolism in TLR3-activated RPE (Figure 1C) and this was accompanied by increased glycolytic reserve post-oligomycin (Figure S1A and S1B).

**Figure 1.**
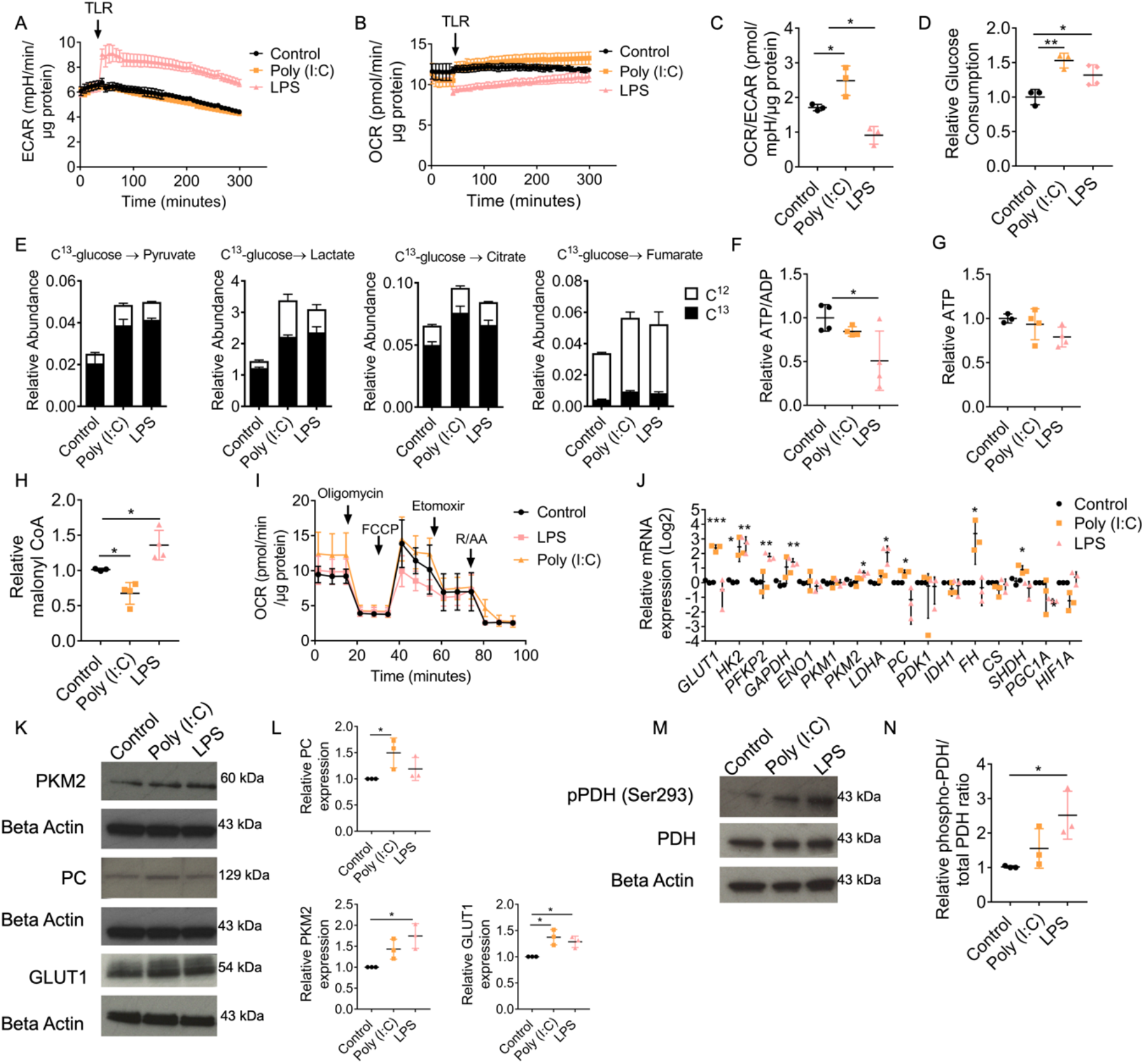
RPE has a differential metabolic response to varying TLR agonists. (A) Real-time changes in ECAR following injection of LPS (1μg/ml) or Poly (I:C) (10μg/ml) to ARPE-19 (*n=3*). (B) Real-time changes in OCR following injection of LPS (1μg/ml) or Poly (I:C) (10μg/ml) to ARPE-19 (*n=3*). (C) Relative OCR/ECAR ratio following treatment with LPS (1μg/ml; 30mins) or Poly (I:C) (10μg/ml; 6h) to ARPE-19 (*n=3*). (D) ARPE-19 treated for 24h with either LPS (1μg/ml) or Poly (I:C) (10μg/ml); glucose concentrations were measured in the media prior/following treatment and expressed as relative consumption (*n=3*). (E) Uniformly labelled C13-glucose incorporation into ARPE-19 metabolites treated for 24h with either LPS (1μg/ml) or Poly (I:C) (10μg/ml); relative abundance of C13 and C12 including pyruvate, lactate citrate and fumarate (*n=3*). ATP/ADP ratio (F) and relative ATP levels (G) of ARPE-19 stimulated for 24h with either LPS (1μg/ml) or Poly (I:C) (10μg/ml) (*n=4*). (H) Relative levels of malonyl CoA detected in whole cell lysates of ARPE-19 stimulated with either LPS (1μg/ml) or Poly (I:C) (10μg/ml) for 24h (*n=3*). (I) Modified mitochondrial stress test including a third injection of etomoxir (3μg/ml) of ARPE-19 stimulated with either 30min LPS (1μg/ml) or 6h Poly (I:C) (10μg/ml) (*n=3*). (J) ARPE-19 treated for 24h with either LPS (1μg/ml) or Poly (I:C) (10μg/ml); RNA was extracted and converted to cDNA; RT-PCR was used to determine the relative gene expression of targets involved in glycolysis or the TCA cycle (*n=3*). (K) ARPE-19 were treated with either LPS (1μg/ml) or Poly (I:C) (10μg/ml) for 24h; protein was extracted and immunoblot analysis was used to determine the expression of PKM2, GLUT1 and PC. (L) Quantification of immunoblots presented in (K) (*n=3*). (M) ARPE-19 treated with LPS (1μg/ml) for 30min or Poly (I:C) (10μg/ml) for 24h; protein was extracted and immunoblot analysis was used to determine the phosphorylation of pyruvate dehydrogenase. Quantification of immunoblots presented in (M) (*n=3*). Data are expressed as means ± SD from at least three independent experiments. (A-C and I) Represents the biological repeats from three independent experiments (*n=3*); each biological repeat is the mean of two technical repeats (two seahorse wells per experiment). One-way ANOVA with Dunnet’s multiple comparisons test; *p<0.05, **p<0.01, ***p<0.001.

To further confirm the differential bioenergetic adaptations of the RPE to TLR stimulation, abundance of intracellular metabolites stable isotope tracer analysis (SITA) was conducted with [U-13C]-glucose (schematic in Fig. S2A). SITA indicated that treatment with both LPS and Poly (I:C) increased the incorporation of C13 glucose into glyceraldehyde-3-phosphate (G3P) (Figure S2B), pyruvate and lactate (Figure 1E), indicating increased glycolytic flux. Both treatments led to an increased incorporation of C13 glucose into TCA metabolites citrate and fumarate (Figure 1E) and succinate (Figure S2C), aspartate (Figure S2E) and glutamate (Figure S2G). The increased labelled succinate and aspartate observed was attributed to an increase in the M+2 mass isotopologues (Figure S2D and S2F). Poly (I:C) treatment increased M+4 glutamate abundance (Figure S2H), indicative of TCA cycling. Incorporation into the TCA intermediates was more pronounced with Poly (I:C) treatment than LPS, confirming increased mitochondrial activity. Furthermore, the relative abundance of unlabelled C12 intermediates of fumarate, succinate, aspartate and glutamate substantially increased with Poly (I:C) treatment, suggesting other fuels may contribute to these metabolite pools (Figure 1E, Figure S2C and Figure S2G). Our data indicate increased glycolysis in LPS stimulated ARPE-19 cells (Figure 1C) and this was confirmed by an increase in glucose consumption (Figure 1D). We further demonstrated that LPS treatment (but not Poly (I:C)) decreased the ATP/ADP ratio in ARPE-19 cells (Figure 1F), indicating a switch away from mitochondrial metabolism. We also found that there are no significant alterations to ATP turnover in these cells following LPS treatment for 24h (Figure 1G).

In addition to glucose-derived carbon entering the TCA cycle, we investigated how fatty acid metabolism was affected by TLR-signalling. First, we measured malonyl CoA levels in treated cells to assess fatty acid synthesis. We found that Poly (I:C) stimulation led to a reduction of malonyl CoA, whereas LPS stimulation increased malonyl CoA levels (Figure 1H). we have explored alterations to fatty acid oxidation through a modified mitochondrial stress test, including the third injection of etomoxir. We have found that in response to LPS fatty acid oxidation is reduced in RPE cells, whereas with Poly (I:C) treatment, fatty acid oxidation is increased (Figure 1I).

Consistent with increased rates of glycolysis, RT-PCR analysis demonstrated greater expression of glycolytic enzymes following LPS treatment (Figure 1J). Poly (I:C) treatment also increased expression of glycolytic enzymes, GLUT1 and TCA enzymes (Figures 1J). To confirm that the changes of gene expression were consistent with protein expression, immunoblotting was performed on three targets with significant alterations following TLR stimulation (GLUT1, pyruvate kinase M2 (PKM2), and pyruvate carboxylase (PC), yielding comparable results (Figure 1K-L).

The pyruvate dehydrogenase (PDH) is inactivated by the phosphorylation on a highly conserved serine residue (Ser293) in the E1 subunit inhibits activity [19], restricting carbon entry into the TCA cycle. Consistent with the increased abundance of pyruvate and lactate, we observed that LPS treatment increased the phosphorylation, and thus inactivation of PDH (Figure 1M-N). Furthermore, inhibition of upstream kinase 1 (PDK1) reverses the LPS-mediated effects on ECAR and OCR (Figures S1C-F). Collectively, these data show that LPS stimulation of RPE drives a rapid switch to aerobic glycolysis, whereas Poly (I:C) stimulation increases oxidative glucose metabolism and TCA activity.

We next investigated the structural changes to mitochondria which occur under innate immune activation. Mitochondria are hubs of metabolic activity, antiviral responses and cell death cascades which continually remodel their structure to facilitate cell processes [20]. Dynamic changes in mitochondrial remodelling are acutely responsive to changes in cell metabolism [21]. Treatment with Poly (I:C), but not LPS increased mitochondrial size relative to unstimulated controls (Figure S3A-B). Additionally, there was shift in mitochondrial morphologies, with decreased fragmented mitochondria, and an increase in long and short tubular mitochondria observed with TLR stimulation (Figure S3F).

Finally, we investigated whether the metabolic changes we observed are applicable to other RPE cells and not just unique to this immortalised RPE cell line, we performed these experiments on primary murine RPE. Similar results were observed (Figure S1G-I). To determine whether the metabolic changes observed in RPE were cell specific, other TLR-expressing cells within the retina (human Müller glia and murine bone-marrow derived mast cells (BMMC)) were used as comparators. We observed a comparable metabolic response in Muller glia, whereas BMMC increased their aerobic glycolysis in response to both agonists (Figure S1J-M).

### Alternate bioenergetic profiles are regulated by AMP-activated kinase

Like immune cells [22], we hypothesised that in the RPE, TLR-induced metabolic responses are governed by AMPK activation status. ARPE-19 cells were cultured in the presence or absence of Poly (I:C), or LPS at a range of time points, from 30 minutes to 24 hours. Treatment of cells with Poly (I:C) had a maximal effect at 24 hours, up-regulating the activity of AMPK, demonstrated by measuring phosphorylation of its downstream target, acetyl-coA carboxylase (ACC) (Figure 2A and 2C). Stimulation with LPS reduced the phosphorylation of both AMPK and ACC observed at 30 minutes (Figure 2B-C), supporting the metabolic changes observed in Figure 1A and B.

**Figure 2.**
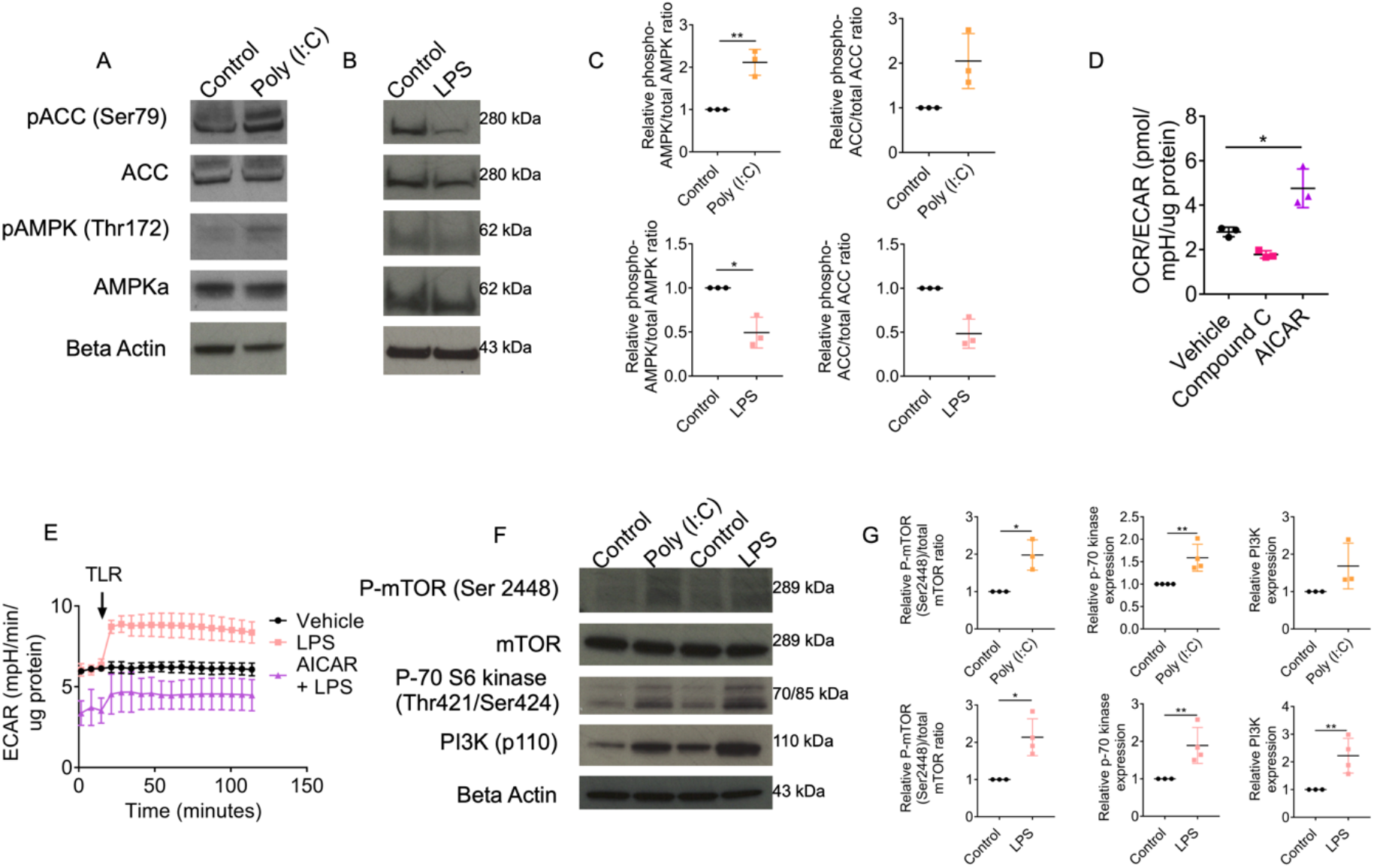
Alternate bioenergetic profiles are regulated by the activity of AMP-activated kinase. (A) ARPE-19 were treated with Poly (I:C) (10μg/ml) for 6h; protein was extracted and immunoblot analysis was used to determine the phosphorylation of AMPK and ACC. (B) ARPE-19 were treated with LPS (1μg/ml) for 30min; protein was extracted and immunoblot analysis was used to determine the phosphorylation of AMPK and ACC. (C) Quantification of immunoblots presented in A and B (*n=3*). (D) Relative basal OCR and ECAR of ARPE-19 treated for 30min with either AICAR (1mM) or compound C (40μM) (*n=3*). (E) Real-time changes in relative ECAR following injection of LPS (1μg/ml) to ARPE-19 cells ± AICAR (1mM) pre-treatment for 30min (*n=3*). (F) ARPE-19 were treated with Poly (I:C) (10μg/ml) for 6h or LPS (1μg/ml) for 1h; protein was extracted and immunoblot analysis was used to determine the activation of mTOR, p-70 s6 kinase and PI3-kinase. (G) Quantification of immunoblots presented in F (*n=3*). (A, B and F) Data are expressed as means ± SD from three independent blots. (D-E) Represents the biological repeats from three independent experiments (*n=3*); each biological repeat is the mean of two technical repeats (two seahorse wells per experiment). (D) One-way ANOVA with Dunnet’s multiple comparaisons test; \**p*<0.05. (E) Unpaired Student’s T-test; \**p*<0.05, \*\**p*<0.01.

Confirming its role as a key metabolic regulator, we find that pharmacological activation and inhibition of AMPK have opposing effects on RPE metabolism (Figure 2D). The increase in OCR/ECAR with AICAR treatment supports the hypothesis that AMPK inhibits aerobic glycolysis [23]. Further to this, we demonstrate that pharmacological activation of AMPK attenuates the switch to aerobic glycolysis in response to LPS (Figure 2E).

The downstream phosphoinositide 3-kinase (PI3K)/ mammalian target of rapamycin (mTOR) signalling pathway was also examined following TLR activation in RPE cells. Poly (I:C) treatment increased mTOR phosphorylation on Ser 2448 of the catalytic subunit and increased the expression of mTOR substrate/surrogate p-70 S6 kinase (Figure 2F-G). Stimulation with LPS upregulated the expression of PI3K p110 (Figure 2F-G). Downstream of PI3K, LPS treatment led to increased activity of mTOR and p-70 S6 kinase (Figure 2F-G).

These data demonstrate that the differential metabolic responses to TLR-stimulation in the RPE are regulated by AMPK activity. The activation of downstream-signalling pathways supports the appropriate alterations of RPE metabolism that facilitate effector functions under stress.

### Interleukin-33 increases mitochondrial metabolism in RPE

We have previously shown that TLR agonists induce the expression of IL-33 in RPE cells, and this regulates cell responses and tissue homeostasis [8]. By extrapolation from observations of the metabolic changes associated with cytokine signaling in other immune cells [24], we hypothesised that activation of RPE through the IL-33/ST2 axis also induces a metabolic change.

Alterations to the OCR/ECAR ratio were observed at 24h of treatment of ARPE-19 cells with IL-33 (Fig. S5C-D), with no effect on RPE viability (Figure S5E and S5F). A mitochondrial stress test identified increases in oxidative metabolism following treatment, particularly increases in spare respiratory capacity (Figure 3A; Figure S5G). Glycolytic metabolism was also increased (Figure 3B; Figure S5H). Primary murine RPE also increased their aerobic metabolism in response to rmIL-33 treatment in an IL-1RL1-dependent manner (Figure S5J).

**Figure 3.**
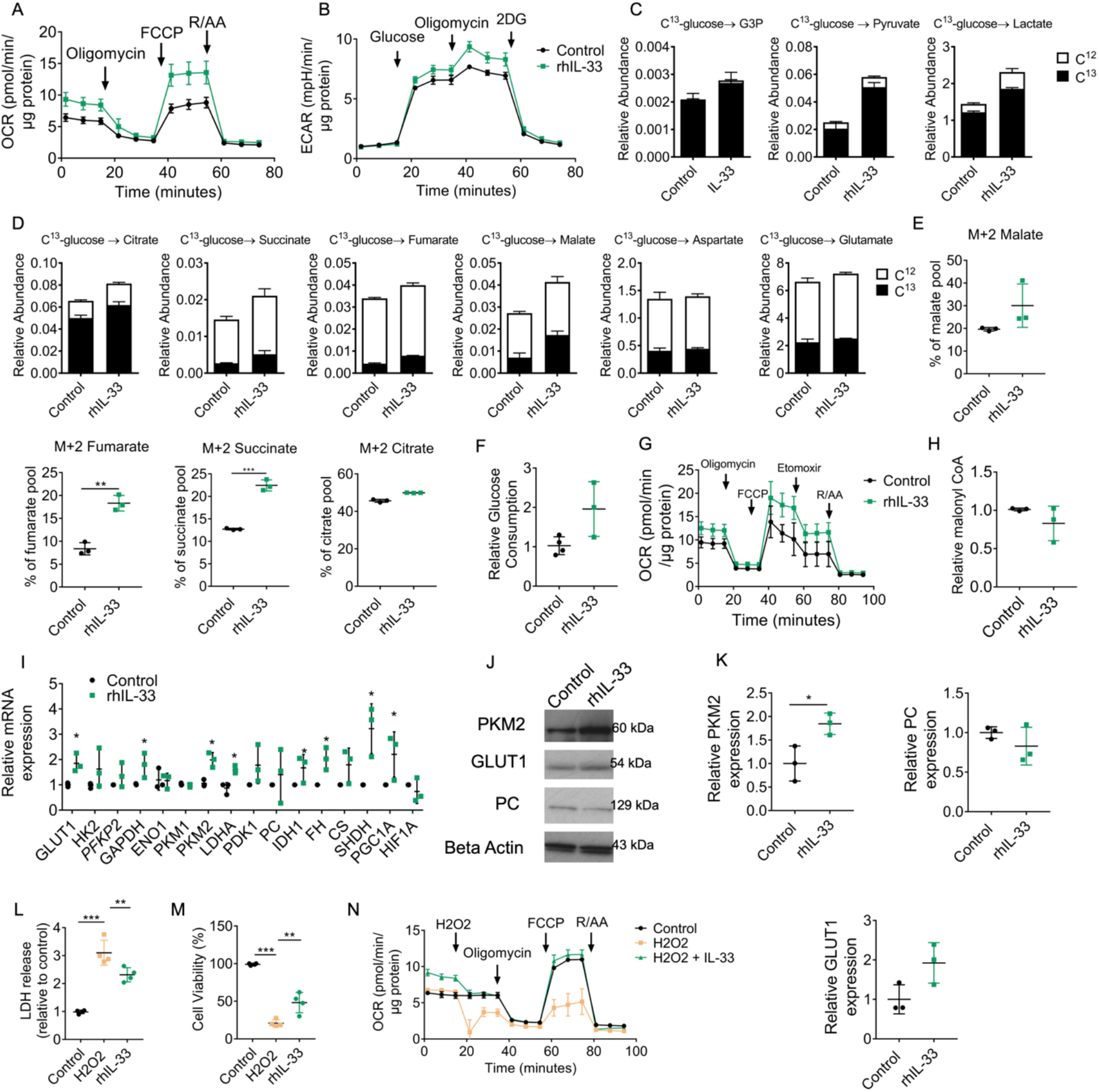
Interleukin-33 increases bioenergetic demand. (A) Mitochondrial stress test from ARPE-19 treated with rhIL-33 (100ng/ml) for 24h; XF injections were oligomycin (1μM), FCCP (0.5μM) and rotenone/antimycin A (1μM) (*n=3*). (B) Glycolysis stress test from ARPE-19 treated with rhIL-33 (100ng/ml) for 24h; XF injections were glucose (10mM), oligomycin (1μM) and 2-deoxyglucose (100mM) (*n=3*). (C) Uniformly labelled C13-glucose incorporation into ARPE-19 glycolytic metabolites treated for 24h with rhIL-33 (100ng/ml); relative abundance of C13 and C12 including 3PG, pyruvate, lactate (*n=3*). (D) Uniformly labelled C13-glucose incorporation into ARPE-19 TCA metabolites treated for 24h with rhIL-33 (100ng/ml); relative abundance of C13 and C12 including malate, fumarate, succinate, citrate, aspartate and glutamate (*n=3*). (E) Mass isotopomer distribution of C13-glucose-derived carbon into fumarate (M+2), succinate (M+2) and citrate (M+2) metabolite pools (*n=3*). (F) ARPE-19 treated for 24h with rhIL-33 (100ng/ml); glucose concentrations were measured in the media prior/following treatment and expressed as relative consumption (*n=3*). (G) Modified mitochondrial stress test including a third injection of etomoxir (3μg/ml) of ARPE-19 cells treated for 24h with rhIL-33 (100ng/ml) (*n=3*). (H) Relative levels of malonyl CoA detected in whole cell lysates of ARPE-19 (*n=3*). (I) ARPE-19 treated for 24h with rhIL-33 (100ng/ml); RNA was extracted and converted to cDNA; RT-PCR was used to determine the relative gene expression of targets involved in glycolysis or the TCA cycle (*n=3*). (J-K) ARPE-19 treated for 24h with rhIL-33 (100ng/ml); protein was extracted and immunoblot analysis was used to determine the expression of PKM2, GLUT1 and PC (*n=3*). (L) ARPE-19 were treated with rhIL-33 (100ng/ml) for 12h before treatment with H2O2 (1mM) for 24h; LDH release was quantified in supernatants and expressed as relative to an untreated control (*n=4*). (M) ARPE-19 were treated with rhIL-33 (100ng/ml) for 12h before treatment with H2O2 (1mM) for 24h; an MTT assay was used to determine cell viability and expressed as a % of an untreated control (*n=4*). (N) Modified mitochondrial stress test of ARPE-19 treated for 24h with rhIL-33 (100ng/ml); XF injections were H2O2 (1mM), oligomycin (1μM), FCCP (0.5μM) and rotenone/antimycin A (1μM) (*n=3*). Data are expressed as means “ SD from at least three independent experiments. (A, B and G) Represents the biological repeats from three independent experiments (*n=3*); each biological repeat is the mean of two technical repeats (two seahorse wells per experiment). (N) Represents the biological repeats from three independent experiments (*n=3*); each biological repeat is either the mean of two technical repeats or a single technical repeat (one or two seahorse wells per experiment). One-way ANOVA with Dunnet$s multiple comparaisons test; *p<0.05,**p<0.01, ***p<0.005.

SITA with [U-13C]-glucose indicated increased C13 incorporation into G3P, pyruvate and lactate pools with IL-33 treatment (Figure 3C). We also observed increased C13 incorporation into TCA metabolites fumarate, malate, succinate, and citrate (Figure 3D). The distribution of the M+2 mass isotopologue was increased in the fumarate, malate and succinate pools, following IL-33 treatment (Figure 3E). Glucose consumption was not significantly affected with Il-33 treatment (Figure 3F).

In addition to glucose-derived carbon entering the TCA cycle, we investigated how fatty acid metabolism was affected by IL-33 signaling. Consistent with the observed activation of AMPK/ inactivation of ACC (Figure S7A-C), we observed an increase in fatty acid oxidation (Figure 3G). No changes were observed to the levels of malonyl coA with IL-33 treatment (Figure 3H). RT-PCR analysis further supported an increase in gene expression of glycolytic and TCA enzymes upon IL-33 stimulation (Figure 3I). Protein expression of PKM2, GLUT1 and PC correlated with gene expression analysis (Figure 3J-K).

Given the functional significance of IL-33/ST2 signalling on mitochondrial capacity, we considered whether this pathway protects against oxidative damage to the RPE. To this end, H2O2 was used as an in vitro platform for inducing reactive oxygen species within the RPE [25]. A dose-dependent relationship was observed between H2O2 concentration and cell viability (Figure S6A). RPE cells treated with H2O2 for 24h exhibited reduced mitochondrial function and glycolytic capacity (Figure S6B-C). Pre-treatment with IL-33 conferred some protection against H2O2-induced RPE dysfunction. This was indicated by reduced LDH release, an increase in cell viability to H2O2 alone (Figure 3L-M) and maintenance of mitochondrial OXPHOS both at 12 and 24h (Figure 3N; Figure S6D). IL-33 was unable to rescue the H2O2-mediated defects in glycolytic metabolism with 12h of treatment (Figure S6E). Treatment with IL-33 for 24h had no significant initial !rescuing” effect to ECAR post-H2O2, but at the greatest dose (100ng/ml) ECAR levels were restored to controls by the end of the assay (Figure S6F). When IL-33 was added at the same time as H2O2 there was no protective effect (Figure S6G), likely due to oxidation and consequent inactivation of IL-33 [26]. We could not attribute the effect of IL-33 treatment to expression changes in antioxidant enzymes or pro-survival proteins (Figure S7D-J). We observed that IL-33 induced a shift in mitochondrial morphology towards a more elongated phenotype and increased mitochondrial size (Figure S8). Despite the observation that IL-33 increases PGC1A, we find that IL-33 treatment has no significant effects on relative mtDNA expression levels (Figure S8F) and therefore suggests that biogenesis may be unaffected (Figure S8F).

### IL-33 is essential for the maintenance of mitochondrial respiration in RPE

Recent studies have determined a novel role for IL-33 in the maintenance of mitochondrial respiration in beige and brown adipocytes [18]. This coupled with the data we have shown thus far, led us to further examine the effect of IL-33 loss on RPE bioenergetics. To knockdown (KD) IL-33 in ARPE-19 cells, four preselected siRNA duplexes, each targeting different sequences of the human IL33 gene, were utilized. The effectiveness of siRNA KD was confirmed both at the protein and mRNA level (Figure 4A-B). Next, we examined the bioenergetic status using extra-cellular flux analysis. Mitochondrial stress analysis identified reduced maximal respiration compared to the scrambled siRNA, while no changes were observed in basal respiration. (Figure 4C-E). A glycolysis stress test identified increased glycolysis following IL-33 KD (Figure 4F-G). The increased glycolysis observed with IL-33 depletion was further confirmed by the increased mRNA expression of key glycolytic enzymes LDHA and GAPDH, and a decrease in the key TCA enzyme PC (Figure 4H). Gene expression correlated to the protein expression of targets PKM2, GLUT1 and PC (Figure 4I-J). The decreased OCR/ECAR ratio confirmed the increased aerobic glycolysis compared to mitochondrial activity (Figure 9A). The addition of rhIL-33 to the IL-33 KD cells was unable to recompense impaired mitochondrial function significantly (Figure 4K), indicating that it is mainly the intrinsic IL-33 that regulates the metabolic responses we observed.

**Figure 4.**
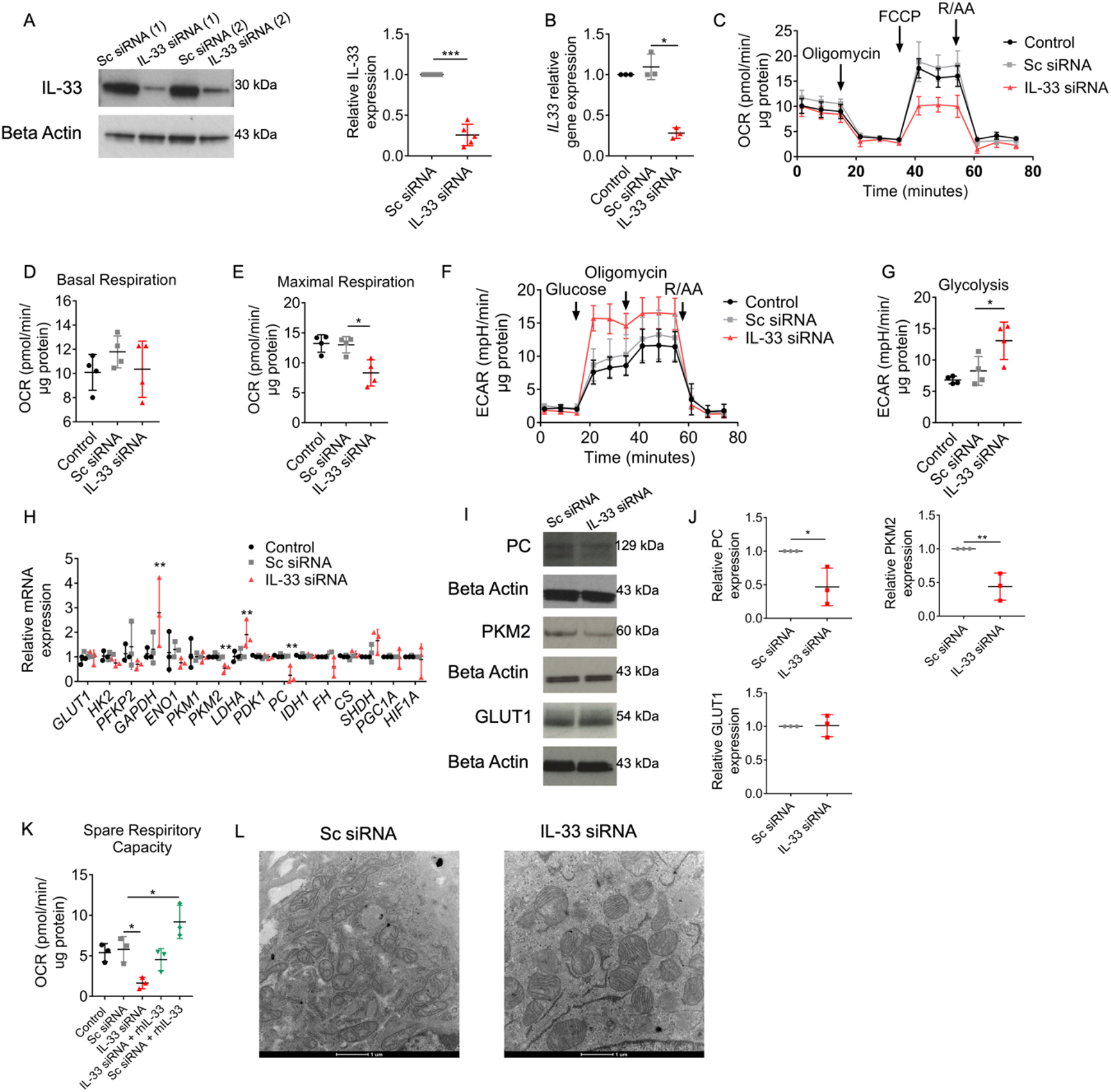
IL-33 is essential for the maintenance of mitochondrial respiration in RPE. (A) Immunoblot of IL-33 following transfection of ARPE-19 with IL-33 siRNA or scrambled siRNA (*n=5*). (B) Gene expression of IL33 following transfection of ARPE-19 with IL-33 siRNA or scrambled siRNA (*n=3*). (C) Mitochondrial stress test following transfection of ARPE-19 with IL-33 siRNA or scrambled siRNA; XF injections were oligomycin (1μM), FCCP (0.5μM) and rotenone/antimycin A (1μM) (*n=4*). (D-E) Parameters calculated from (C) (as detailed in methods) (*n=4*). (F) Representative glycolysis stress test following transfection of ARPE-19 with IL-33 siRNA or scrambled siRNA; XF injections were glucose (10mM), oligomycin (1μM) and 2-deoxyglucose (100mM) (*n=4*). (G) Parameter calculated from (F) (as detailed in methods) (*n=4*). (H) ARPE-19 were transfected with IL-33 siRNA or scrambled siRNA; RT-PCR was used to determine the relative gene expression of targets involved in glycolysis or the TCA cycle (*n=3*). (I-J) ARPE-19 cells were transfected with either IL-33 siRNA or scrambled siRNA; protein was extracted and immunoblot analysis was used to determine the expression of PKM2, GLUT1 and PC (*n=3*). (K) Following transfection of ARPE-19 with IL-33 siRNA or scrambled siRNA, cells were treated with rhIL-33 for 24h; spare respiratory capacity was analysed using a mitochondrial stress test; XF injections were oligomycin (1μM), FCCP (0.5μM) and rote-none/antimycin A (1μM) (*n=3*). (L) Representative transmission electron microscopy of ARPE-19 cells transfected with IL-33 siRNA or scrambled siRNA; Magnification 4500x. Data are expressed as means” SD from at least three independent experiments. (C-G) Represents the biological repeats from three independent experiments (*n=3*); each biological repeat is the mean of two technical repeats (two seahorse wells per experiment). (K) Represents the biological repeats from three independent experiments (*n=3*); each biological repeat is either the mean of two technical repeats or a single technical repeat (one or two seahorse wells per experiment). (A) Unpaired Student$s T-test; \**p*<0.05. (B, E, G, H, I, J and K) One-way ANOVA with Dunnet$s multiple comparaisons test; \**p*<0.05, \*\**p*<0.01.

Given the significant reduction in mitochondrial metabolism in the absence of IL-33, we utilised transmission electron microscopy to identify any visible structural changes in IL-33KD RPE mitochondria. Compared to a scrambled siRNA, IL-33KD mitochondria appeared to be larger and more fragmented (Figure 4L). Image analysis of mitochondrial samples showed that mitochondrial diameter and area was increased with IL-33 siRNA and confirmed that a greater percentage of mitochondria were of a fragmented phenotype (Figure S9B). The fragmented mitochondrial phenotype observed in the IL-33KD group was reminiscent of previously published data in IL-15 T-memory cells, whereby mitochondrial fission promoted an increased glycolytic capacity [27].

### Bioenergetic analysis of IL-33−/− primary RPE

To investigate further the role of intrinsic IL-33, we performed a bioenergetic analysis of primary RPE derived from Il33−/− mice. Using a mitochondrial stress test, it was observed that in comparison to WT, Il33−/− mice had reduced maximal respiration and spare respiratory capacity (Figure 5A and 5B). No significant changes were observed in basal respiration or ATP-production (Figure 5A). Like IL-33 KD ARPE-19 cells, we observed that primary Il33−/− RPE had increased glycolysis, and yet lacked glycolytic reserve measured post-oligomycin treatment (Figure 5C and 5D). There was no significant effect on the OCR/ECAR ratio (Figure 5E). Together, these results suggest that in the absence of IL-33 the relative contribution of glycolysis to ATP production compared with OXPHOS is greater. The addition of rmIL-33 to primary Il33−/− RPE was unable to recompense impaired mitochondrial function (Figure 5F and 5G).

**Figure 5.**
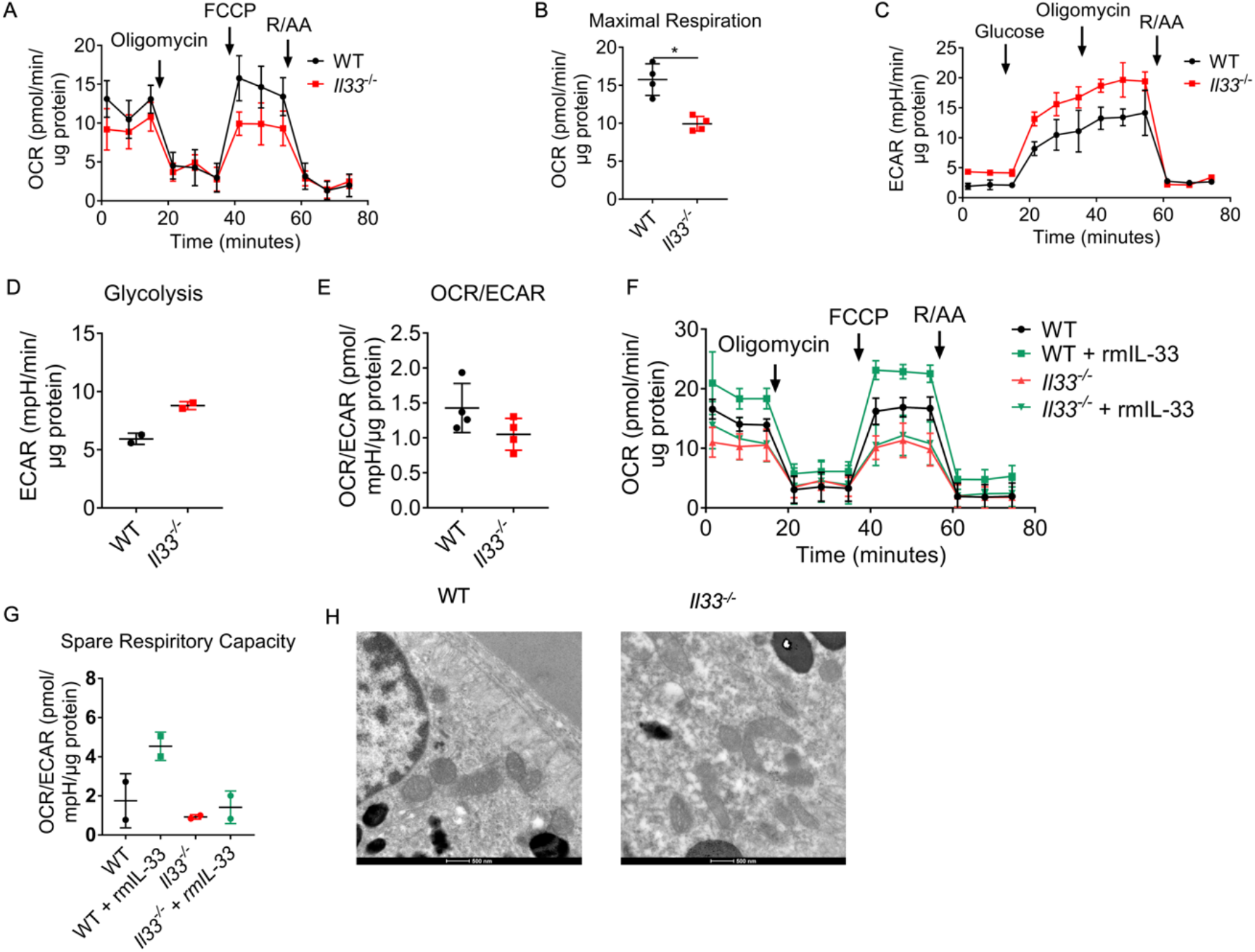
Bioenergetic analysis of Il-33−/− primary RPE. (A) Mitochondrial stress test of WT and Il33−/− primary murine RPE; XF injections were oligomycin (1μM), FCCP (0.5μM) and rotenone/antimycin A (1μM) (*n=4*). (B) Parameters calculated from (A) (as detailed in results) (*n=4*). (C) Glycolysis stress test of WT and Il33−/− primary murine RPE; XF injections were glucose (10mM), oligomycin (1μM) and 2-deoxyglucose (100mM) (*n=2*). (D) Parameters calculated from (C) (as detailed in methods) (*n=2*). (E) OCR/ECAR ratio of WT and Il33−/− primary murine RPE (*n=4*). (F) Primary murine RPE were isolated from mice both WT and Il33−/− mice and treated for 24h with rmIL-33; a mitochondrial stress test was used to assess mitochondrial function; XF injections were oligomycin (1μM), FCCP (0.5μM) and rotenone/antimycin A (1μM) (*n=2*). (G) Parameters calculated from (F) (as detailed in methods) (*n=4*). (H) Representative transmission electron microscopy of RPE from WT and Il33−/− mice. Magnification 9300x. (A, B and E) Represents the biological repeats from four independent experiments (*n=4*); each biological repeat is either the mean of two technical repeats or a single technical repeat (one or two seahorse wells per experiment). (C-D, F-G) Represents the biological repeats from two independent experiments (*n=2*); each biological repeat is either the mean of two technical repeats or a single technical repeat (one or two seahorse wells per experiment). (A, B and E) Represents data from eight eyes (four mice) per group. (C-D, F-G) Represents data from four eyes (two mice) per group. (A-B) Unpaired Student$s T-test; *p<0.05, **p<0.01. (F-G) One-way ANOVA with Dunnet$s multiple comparisons test; \**p*<0.05, \*\**p*<0.01.

Further supporting a nuclear role for IL-33, we observed that primary RPE derived from Il1rl1−/− mice lacked any metabolic phenotype (Figure S10). This excluded the possibility that metabolic alterations could have been attributed to lack of IL-33 release and autocrine/paracrine signalling through cognate receptor ST2.

We utilised transmission electron microscopy to identify any visible structural changes in Il33−/− RPE mitochondria. Compared to WT, Il33−/− mitochondria appeared to be smaller and more irregular in size, (Figure 5H). Image analysis of mitochondrial samples showed that mitochondrial diameter was decreased in Il33−/− (Figure S11A). No significant changes were observed in mitochondrial area, but there was increased mitochondrial number/field in the Il33−/− group (Figure S11B and S11C). In contrast to the data presented from IL-33 loss in human cells, image analysis identified an increase in !short tubular” mitochondria in the Il33−/− group (Figure S11D).

### Role of nuclear IL-33 in mitochondrial metabolism

Following the identification of metabolic perturbations associated with IL-33 absence in vitro and ex vivo, we examined whether increased IL-33 expression in the RPE would have an adverse impact on cell metabolism. Using a CRISPRcas9 activation plasmid, the expression of the human IL33 gene was upregulated in ARPE-19 cells. The effectiveness of CRISPRcas9-mediated over-expression of IL-33 was confirmed both at the RNA and protein level in whole cell lysates (Figure 6A and 6B). As IL-33 functions as a dual function cytokine, residing both within the nucleus and acting extracellularly [10], it was necessary to identify the subcellular location of IL-33 following overexpression. Increased IL-33 was observed in the nuclear subcellular fraction (Figure 6C). A mitochondrial stress test was used to identify changes in OXPHOS parameters associated with IL-33 overexpression and this showed increased maximal (FCCP-induced) respiration rates (Figure 6D and 6E). Glucose-starved cells were subjected to a glycolysis stress test to identify changes in glycolytic parameters and IL-33 overexpression was found to significantly increase glycolytic metabolism (Figure 6F and 6G). This was accompanied by increased gene expression of glycolytic and TCA cycle enzymes (Figure 6H). Protein expression of GLUT1, PC and PKM2 was confirmed at the protein level (Figure 6I-J). We finally confirmed alterations to mitochondrial function were accompanied by structural alterations. Image analysis indicated that mitochondria in the IL-33 overexpression group formed long elongated tubules (Figure 6K; Figure S12), a structural phenotype associated with increased OXPHOS in T-cells [27].

**Figure 6.**
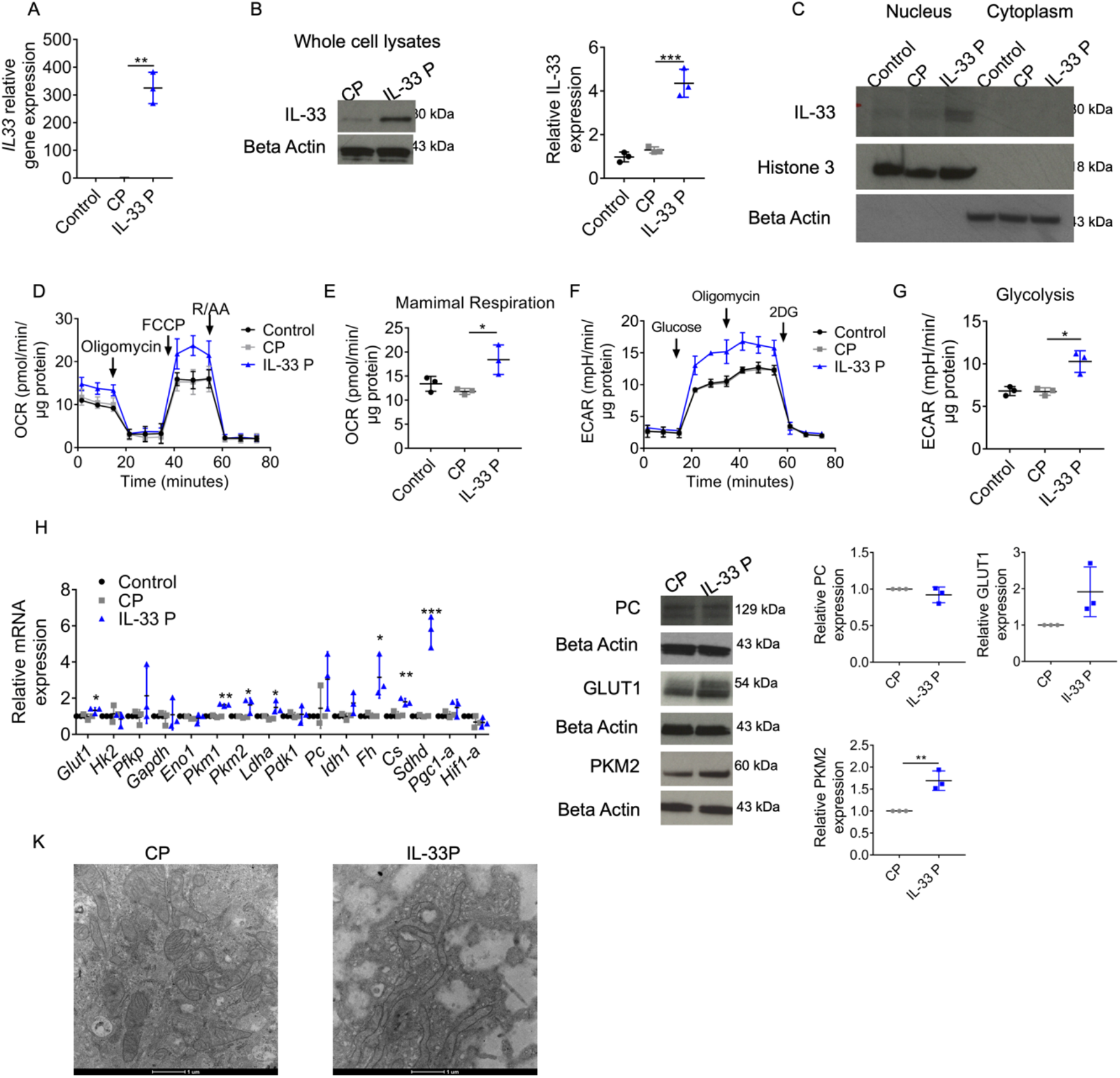
Role of nuclear IL-33 in mitochondrial metabolism. (A) Gene expression of IL33 following transfection of ARPE-19 with either an IL-33 activation plasmid or scrambled gRNA activation plasmid (*n=3*). (B) Representative immunoblot of IL-33 expression in ARPE-19 transfected with either an IL-33 activation plasmid or scrambled gRNA activation plasmid (*n=3*). (C) ARPE-19 were transfected with either an IL-33 activation plasmid or scrambled gRNA activation plasmid; cell lysates were split into nuclear or cytoplasmic fractions and western blotting was used to determine the subcellular location of IL-33 (*n=2*). (D) Mitochondrial stress test following transfection of ARPE-19 with either an IL-33 activation plasmid or scrambled gRNA activation plasmid; XF injections were oligomycin (1μM), FCCP (0.5μM) and rote-none/antimycin A (1μM) (*n=3*). (E) Parameters calculated from (D) (as detailed in methods) (*n=3*). (F) Glycolysis stress test following transfection of ARPE-19 with either an IL-33 activation plasmid or scrambled gRNA activation plasmid; XF injections were oligomycin (1μM), FCCP (0.5μM) and rotenone/antimycin A (1μM) (*n=3*). (G) Parameters calculated from (F) (as detailed in methods) (*n=3*). (H) ARPE-19 were transfected either an IL-33 activation plasmid or scrambled gRNA activation plasmid; RT-PCR was used to determine the relative gene expression of targets involved in glycolysis or the TCA cycle (*n=3*). (I-J) ARPE-19 were transfected either an IL-33 activation plasmid or scrambled gRNA activation plasmid; protein was extracted and immunoblot analysis was used to determine the expression of PKM2, GLUT1 and PC (*n=3*). (K) Representative transmission electron microscopy of ARPE-19 cells transfected with an IL-33 activation plasmid or scrambled gRNA activation plasmid. Magnification 4500x. Data are expressed as means " SD from at least three independent experiments. (D-G) Represents the biological repeats from three independent experiments (*n=3*); each biological repeat is the mean of two technical repeats (two seahorse wells per experiment). (C) Represents two independent immunoblots. One-way ANOVA with Dunnet$s multiple comparisons test; *p<0.05, **p<0.01, ***p<0.001.

### Nuclear IL-33 promotes oxidative glucose metabolism

Since we observed an increase in ECAR (Figure 4F-G) and extracellular lactate (Figure S13A). in the absence of IL-33, we investigated whether IL-33 might regulated pyruvate import into the TCA. Pyruvate enters the mitochondria through the mitochondrial pyruvate carrier complex (MPC) consisting of components MPC1/2 [28]. Overexpression of IL-33 led to increased MPC1 and MPC1 but had no significant effect on MPC2 and MPC2 expression (Figure 7A and 7B). IL-33 KD reduced the expression of both MPC components at both the gene and protein level (Figure 7C and 7D). Hence, we performed a modified mitochondrial stress test with the additional injection of the MPC inhibitor UK5099 to assess the contribution of aerobic glucose metabolism to total OCR. In ARPE-19 cells with IL-33 overexpression, the increased maximal respiration was largely due to increased pyruvate-dependent respiration (Fig. 7E-F). When a similar experiment was performed on IL-33 siRNA cells it was observed that pyruvate-dependent respiration had a significantly reduced contribution to maximal OCR (Figure 7G-H).

**Figure 7.**
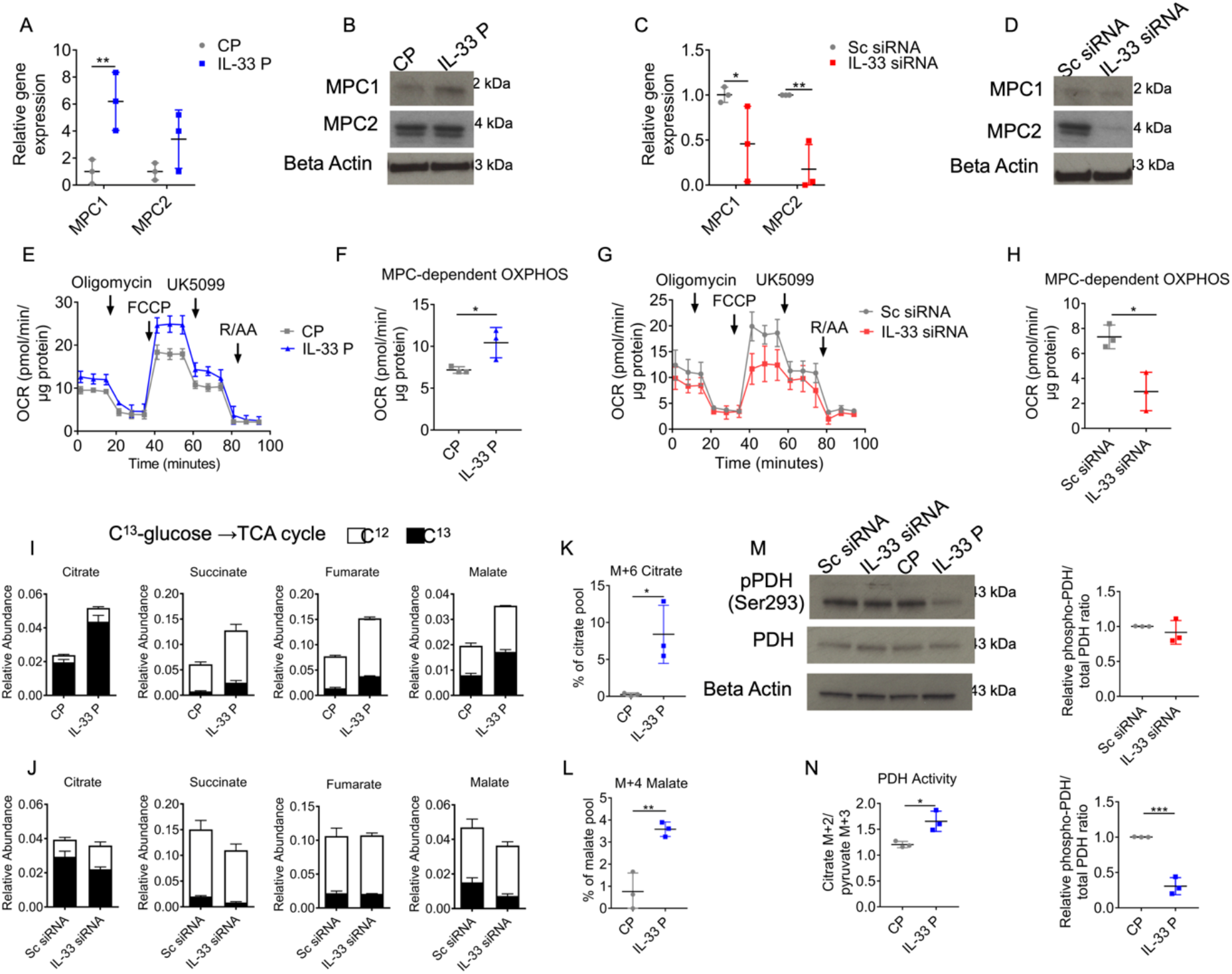
Nuclear IL-33 promotes oxidative glucose metabolism. (A-B) ARPE-19 were transfected with either an IL-33 activation plasmid or scrambled gRNA activation plasmid; (A) RNA was extracted, and RT-PCR was used to determine the expression of *MPC1* and *MPC2* (*n=3*); (B) protein was extracted and western blot analysis was used to determine the expression of MPC1 and MPC2 (*n=3*). (C-D) ARPE-19 were transfected with either an IL-33 siRNA or scrambled siRNA; (C) RNA was extracted, and RT-PCR was used to determine the expression of *MPC1* and *MPC2* (*n=3*); (D) protein was extracted and western blot analysis was used to determine the expression of MPC1 and MPC2 (*n=3*). (E) Modified mitochondrial stress test following transfection of ARPE-19 with either an IL-33 activation plasmid or scrambled gRNA activation plasmid; XF injections were oligomycin (1μM), FCCP (0.5μM), UK5099 (5μM) and rotenone/antimycin A (1μM) (*n=3*). (F) Parameter calculated from (E) (as detailed in methods) (*n=3*). (G) Modified mitochondrial stress test following transfection of ARPE-19 with either an IL-33 siRNA or scrambled siRNA; XF injections were oligomycin (1μM), FCCP (0.5μM), UK5099 (5μM) and rotenone/antimycin A (1μM) (*n=3*). (H) Parameter calculated from (G) (as detailed in methods) (*n=3*). (I) Uniformly labelled C13-glucose incorporation into ARPE-19 TCA cycle metabolites following transfection with either an IL-33 activation plasmid or a scrambled gRNA activation plasmid control; relative abundance of C13 and C12 including succinate, fumarate, malate and citrate (*n=3*). (J) Uniformly labelled C13-glucose incorporation into ARPE-19 TCA cycle metabolites following transfection with either an IL-33 siRNA or scrambled siRNA control (*n=3*). Relative abundance of C13 and C12 including succinate, fumarate, malate and citrate (*n=3*). (K-L) Mass isotopologue distributions (MID) of ARPE-19 TCA cycle intermediates (K) M+6 citrate and (L) M+4 malate, following transfection with either an IL-33 activation plasmid or a scrambled gRNA activation plasmid (*n=3*). (M) ARPE-19 were transfected with either an IL-33 activation plasmid/ scrambled gRNA activation plasmid or an IL-33 siRNA/ scrambled siRNA; western blot analysis was used to determine the phosphorylation status of pyruvate dehydrogenase (*n=3*). (N) Citrate M+3/pyruvate M+3 ratio in ARPE-19 transfected with either an IL-33 activation plasmid or scrambled gRNA activation plasmid (*n=3*). Data are expressed as means " SD from at least three independent experiments. (E-H) Represents the biological repeats from three independent experiments (*n=3*); each biological repeat is the mean of three technical repeats (three seahorse wells per experiment). Unpaired Student$s T-test; \**p*<0.05, \*\**p*<0.01, \*\*\**p*<0.001.

To assess if pyruvate metabolism was the only component affected by the altered expression of IL-33, or if other metabolic pathways feeding into the TCA cycle were involved, a similar experiment was conducted to assess FAO. Etomoxir treatment post mitochondrial uncoupling significantly reduced the OCR, however between control plasmid and IL-33 plasmid groups, this reduction was not significant (Figure S13D-E). With IL-33 loss, we observed that FAO was significantly increased (Figure S13F-G), suggesting that FAO is upregulated in the absence of IL-33 to compensate for defects in pyruvate metabolism.

SITA with [U-13C]-glucose was conducted to further assess how IL-33 altered glucose metabolism in RPE. In ARPE-19 cells with IL-33 overexpression, C13 labelling indicated increased glycolytic flux (Figure S14A). No significant changes were observed in glycolysis !end-point” metabolites pyruvate or lactate (Figure S14A); however, significant increases in C13 enrichment were observed in TCA metabolites (Fig. 7I), indicating that glucose-derived TCA cycle activity was up-regulated with IL-33 overexpression. In contrast, ARPE-19 cells with IL-33 KD had a significant increase in the abundance of C13 lactate (Figure S14B, and decreased C13 enrichment in malate, citrate and succinate (Figure 7J).

Overexpression of IL-33 lead to an increase in the fully labelled citrate mass isotopologue (M+6) (Figure 7K). This pattern will occur in citrate when both oxaloacetate (OAA) and malate are derived from glucose. Increased M+4 aspartate (Figure S14C) and malate (Figure 7L) indicate the increased TCA cycling which occurs with IL-33 overexpression. Labelling patterns using C13-labelled glucose then highlighted differential flux from glycolysis and pyruvate input into the TCA cycle. Glucose-derived pyruvate can enter the TCA cycle through either PDH or PC (Figure S14D). The citrate M+2/pyruvate M+3 ratio can serve as a surrogate for PDH activity, while the citrate M+3/pyruvate M+3 ratio is used as a surrogate of PC activity [29]. IL-33 KD significantly reduced the citrate M+3/pyruvate M+3 (Figure S14F) ratios suggesting a decrease in the activity of the PC complex. Although no significant changes were observed in the citrate M+3/pyruvate M+3 ratio with IL-33 overexpression (Figure S14G), there was a significant increase in the citrate M+2/pyruvate M+3 ratio (Figure 7N), suggesting that PDH activity was augmented with IL-33 plasmid treatment. Increased PDH activity was confirmed by reduced PDH E1 phosphorylation status (Fig. 7M).

Glutamate labelling was unaffected by IL-33 overexpression (Figure S15A-B). However, we show that IL-33 knockdown reduces derived C13 labelling of the M+2 mass isotopologue in glutamate pools (Figure S15C-D). We find no observable C13 labelling detected in α-ketoglutarate pools (Figure S15E-H). The decrease in unlabelled α-ketoglutarate observed with IL-33 overexpression (Figure S15E) suggests that the glutamine metabolism is likely reduced as increased glucose-derived carbon is used to support the TCA cycle. The increase in unlabelled glutamate and α-ketoglutarate (Figure S15G) suggest that glutamine-derived carbon may support the TCA cycle when glucose metabolism is impaired.

Taken together these results indicate that nuclear IL-33 is a critical regulator of pyruvate oxidative metabolism in the RPE. When over expressed, there is increased glycolytic flux into the TCA cycle most likely through increased MPC and PDH activity. The absence of IL-33 reduces the oxidative catabolism of glucose, and as pyruvate is !redirected to lactate, FAO appears to support the TCA cycle.

## Discussion

All cells must be able to maintain their energetic resources to survive when quiescent and upon stress. The RPE provides an excellent model system to study basic cellular metabolism due to its active mitochondrial capacity which maintains retinal homeostasis. RPE stress and metabolic alterations including mitochondrial dysfunction have been demonstrated to be implicated in retinal diseases [6, 30]. However, the causal link between stress, metabolic alterations and retinal degeneration remains unclear. In this study we demonstrate that in response to stress there are intrinsic cellular immune responses dependent on IL-33 that act as a checkpoint for metabolic regulation. Control of this metabolic checkpoint means that the RPE can regulate metabolism for rapid energy production whilst ensuring the maintenance of mitochondrial health. We show here that the innate immune activation of the RPE through TLR signalling is supported by a distinct metabolic profile which is dependent on AMPK. We find that IL-33 signalling leads to broad transcriptional changes in metabolic genes which leads to increased metabolic flux through glycolysis and the TCA cycle. Moreover, we provide evidence for a novel role of intracellular IL-33 as a regulator of RPE metabolism. IL-33 loss increases aerobic glycolysis at the expense of oxidative glucose catabolism. Cells overexpressing IL-33 display increased expression of MPC1 while activating pyruvate dehydrogenase (through dephosphorylation) to facilitate increased pyruvate flux into the TCA cycle.

The concept of a metabolic ‘retinal ecosystem’ was recently introduced by Hurley et al [3]. They propose that energy homeostasis in the retina and RPE relies on a complex and specialized metabolic interplay between distinct cells which has implications for retinal diseases [3]. This study suggests that dysfunctional mitochondrial activity, acquired with age, restricts the concentration of glucose able to reach the photoreceptors and consequently reduces the lactate available to the RPE as a fuel source. This increased reliance on glycolysis may upset the metabolic ecosystem and lead to photoreceptor death. Our data support the proposal that AMPK, a central mediator of the bioenergetic response to innate immune stress, maintains energetic homeostasis in the RPE by increasing mitochondrial ATP production. We observed that a stressor; LPS stimulation of the RPE, inactivates AMPK. This elicits a metabolic shift towards aerobic glycolysis for ATP production, this is consistent with results observed in other immune cells, dendritic cells and macrophages [22, 31]. Interestingly, our data also provide evidence that TLR-3 signalling in the RPE (a mechanism leading to retinal degeneration [32]) is accompanied by the late activation of AMPK and increases in both glycolysis and OXPHOS. These data differ from results observed in dendritic cell populations whereby Poly (I:C) stimulation drives increased aerobic glycolysis and a decrease in OXPHOS [33, 34], highlighting distinct properties of the RPE. Unlike AMPK, mTOR promotes catabolic and proliferative signalling and controls processes such as protein synthesis, metabolism and cell growth that are essential for activated inflammatory cells [35]. The activation of AMPK and mTOR alongside the increases in glycolysis and OXPHOS we observed following TLR stimulation in RPE, supports the concept of a ‘metabolic ecosystem’ essential for photoreceptor and neuronal function.

Mitochondrial DNA (mtDNA) can mediate expression levels of nuclear genes related to complement, inflammation and apoptosis [17]. Studies using cybrid RPE cells with representative mtDNA haplotypes demonstrated that different mtDNA variants exhibit a difference in ATP levels, lactate production and metabolic (glycolytic) enzymes expression, which further dictates altered expression of nuclear encoded genes in complement, innate immunity, apoptosis and proinflammatory signaling pathways [17]. Our previous work demonstrated that the RPE responds to TLR agonists by increasing expression of IL-33. IL-33 is a nuclear cytokine whose expression correlates with mtDNA content in the RPE [8]. Therefore, there are contemporaneous changes in RPE metabolism and expression of IL-33 when mtDNA content changes. We propose that IL-33 acts as a signal in the RPE to adapt or maintain mitochondrial metabolism. Consistent with prior reports on cancer cells [36] and fibroblasts [37] we have identified that IL-33/ST2 signalling is accompanied by increased metabolic (particularly glycolytic) activity. In addition, we found that exogenous IL-33 protects RPE cells against the noxious effects of oxidative stress. IL-33 release is regulated by oxidative stress and antioxidants within the airway epithelium [38]. It is reported to reduce ROS production in fibroblasts [39] and directly enhance the activity of SOD in cardiomyocytes [40]. However, the data presented here demonstrate no change in anti-oxidant enzyme expression but rather an increase in metabolic flux and mitochondrial activity.

Our findings imply that IL-33 is essential for supporting mitochondrial respiration. We found that mice lacking IL-33 have abnormal mitochondrial morphologies. Upon subsequent in vitro investigation we observed that endogenous IL-33 loss increases aerobic glycolysis as identified by increased lactate production from glucose. This increased reliance on glycolysis may therefore upset the metabolic ecosystem and lead to RPE dysfunction. This is consistent with observations in adipose tissue whereby isolated mitochondria from *Il33*−/− mice exhibited profound respiratory defects, including reduced OXPHOS, and enzymatic activity of ETC complexes II and IV [18]. The same study identified an increased expression of genes involved in the catabolism of fatty acids, glucose and amino acids in *Il33*−/− adipose tissue suggesting a similar compensatory mechanism in the face of reduced mitochondrial function [18]. This highlights that IL-33 can have common roles across systems, and our data on its role in retinal metabolism can be extrapolated to other tissues and diseases.

The understanding of IL-33 has developed beyond the primary implications in the induction of T-helper (Th) 2 immune responses to that of a cytokine with a broader activity in Th1 immunity and regulatory responses. In allergic and respiratory diseases, IL-33 activates type 2 innate lymphoid cell (ICL2) that elicit eosinophil recruitment and Th2 differentiation [41, 42]. The pro-inflammatory role is supported by correlation of serum IL-33 with disease severity in atopic dermatitis [43] and through disease attenuation with IL-33 inhibition in murine allergic asthma [44]. IL-33 exacerbates disease progression in rheumatoid arthritis models associated with proinflammatory cytokine production, mast cell degranulation and neutrophil recruitment [45, 46]. Contrary, treatment with IL-33 reduces murine atherosclerosis development [16] and IL-33 polarises macrophages (M2 phenotype) conferring protection against obesity and inflammation whilst improving glucose regulation.

Multiple lines of evidence suggest that increased aerobic glycolysis with IL-33 loss in the RPE occurs at the expense of oxidative glucose catabolism, as IL-33 impairs pyruvate import into the mitochondria through the MPC complex. Genetic support for this was obtained from IL-33 KD RPE, where we found reduced expression of both MPC1 and MPC2 (Fig. 7), which facilitate the transport of pyruvate into the mitochondria [47]. Furthermore, cells overexpressing IL-33 displayed reduced aerobic glycolysis as mitochondrial activity was increased (Fig. 6). The increased expression of MPC complex components and activity of pyruvate dehydrogenase was accompanied by an increased consumption and oxidation of glucose within the TCA cycle. The increased spare respiratory capacity was found to be attributed to pyruvate import following the use of an MPC-specific inhibitor UK5099 [48] (Fig. 7E) We found that IL-33 is a key mediator of transcriptional changes to glycolytic and TCA cycle genes, which either drive the RPE towards aerobic glycolysis or mitochondrial catabolism of pyruvate in its absence or presence, respectively. Altered MPC1/2 expression results in significant metabolic disorders and has been previously shown to contribute to the Warburg effect in cancer cells [49].

While our data have uncovered a role for IL-33 in mitochondrial respiration in RPE, a key question that emerges from our work is how IL-33 regulates expression of MPC1/2 in RPE to facilitate the transport of pyruvate. The protective effect of exogenous recombinant IL-33 on RPE against oxidative stress was associated with increased metabolic flux and mitochondrial activity. In the future, generation of cell-and tissue-specific knockouts of IL-33 will be necessary to pinpoint the critical cell types that release IL-33 and the targets on which it acts to regulate the retinal ‘metabolic ecosystem’.

In conclusion, we have demonstrated that IL-33 constitutes a key metabolic checkpoint that antagonises the Warburg effect to ensure the functional stability of the RPE. The identification of IL-33 as a key regulator of mitochondrial metabolism suggests roles for this cytokine that go beyond its extracellular “alarmin” activities. For example, when RPE is under stress, IL-33 contributes to minimise the effects of oxidative damage to the RPE and bolster mitochondrial metabolism. IL-33 exerts control over mitochondrial respiration in RPE by facilitating pyruvate import into mitochondria via up regulation of MPC expression and may be associated with the capacity of RPE to maintain homeostasis. Therefore, as well as identifying a molecular pathway for activation of mitochondrial respiration in RPE, our results demonstrate that intrinsic cellular IL-33 acts as a metabolic regulator exerting profound effects on retinal metabolism.

## Methods

### Cell lines

Immortalized human retinal pigment epithelium (ARPE-19) and human Müller glial Moorfields/Institute of Ophthalmology-Müller 1 (MIO-M1) cell lines were cultured in DMEM (4.5mg/L) supplemented with 10% heat-inactivated fetal bovine serum, 2mM L-glutamine, 1mM sodium-pyruvate, 0.5μM 2-mercaptoethanol, 100U/ml penicillin, and 100μg/ml streptomycin. All cell lines were maintained at 37°C and were routinely screened for mycoplasma contamination. The human MIO-M1 was purchased from the UCL Business PLC, London, UK. The ARPE-19 cell line is a spontaneously arising cell line derived from human retinal pigment epithelium [ATCC number CRL-2302] [50].

### Primary cells

C57BL/6J mice were purchased from Charles River Laboratories, Margate, UK. All mice were maintained in the animal house facilities of the University of Bristol, according to Home Office Regulations. Animal husbandry and procedures complied with the ARVO (Association for Research in Vision and Ophthalmology) statement for the use of animals in ophthalmic and vision research. IL-33 knockout mice (Il33−/−) were generated as described earlier [51] at the animal house facilities of Trinity College Dublin and sent to the animal house facilities of the University of Bristol when required. IL-1RL1 (ST2) knockout (Il1rl1−/−) C57BL/6 mice (backcrossed for ten generations) were kindly provided by C Emanueli (School of Clinical Sciences, Bristol Heart Institute, University of Bristol) and were generated as described earlier [46].

Primary murine RPE cells were obtained as described earlier [52]. Eyes were enucleated and incubated at 37°C in hyaluronidase for 45min, and in Hank’s balanced salt solution (HBSS) with calcium, magnesium and 10mM HEPES for a further 30min. An incision was made beneath the ora serrata to remove the iris epithelium and cornea. The neural retina and the attachment to the optic nerve were removed. Eyecups were incubated at 37°C in trypsin EDTA for 45min. Eyecups were then transferred into HBSS with 20% FBS and shaken gently to allow the RPE to detach. The RPE sheets were incubated at 37°C in 1ml trypsin EDTA for 1min, 9ml primary RPE media was then added, centrifuged, and supernatant removed. The resulting RPE cells were resuspended in 200μL primary RPE media and homogenously mixed using a p200 pipette. Cells were used within 10 days of extraction. Cells were cultured in alpha-MEM, containing 1% N1 supplement, 1% glutamine-1% penicillin-streptomycin, 1% non-essential amino acid solution, 5% heat-inactivated fetal bovine serum, 20μg/L Hydrocortisone, 250mg/L Taurine and 13ng/L triiodo-thyronin. Purity was assessed by immunoblotting for RPE specific protein RPE65 and rhodopsin contamination from the neurosensory retina.

Bone marrow derived mast cells (BMMCs) were generated from C57BL/6 as previously described [53]. Legs were taken from mice and the bones were cut at the end using angles scissors to expose the red marrow. Using a syringe and a 23-gauge needle DMEM was flushed through the bone. DMEM containing the cell suspension was spun at 14,000g for 5min in a centrifuge cooled to 4°C. The resulting cell pellet was resuspended in DMEM containing IL-3 (10ng/ml) and dispensed into a cell culture flask. The cell suspension was cultured for 5 weeks to allow the generation of a mast cell population. Prior to functional assays, mast cells were sorted into a viable population using a MACS dead cell removal kit and the purity was assessed by FACS based on CD117 and CD45 cell surface marker expression.

### Cell culture

Cells were seeded at a density of 100,000 per well of a 24 well plate and were exposed to different inflammatory stimuli; Poly (I:C) (10μg/ml), LPS (1μg/ml) and IL-33 (100-1ng/ml, as detailed in the results). Oxidative stress was induced by the addition of hydrogen peroxide (1mM). AMPK activity was modulated by activator AICAR (1mM), and inhibitor Compound C (40μM). Unstimulated controls and vehicle controls (DMSO) were always included.

### Western blot

Following treatment, protein extraction was performed using Cell-Lytic-M with the addition of Protease/Phosphatase Inhibitor Cocktail. A commercial nuclear extraction kit was used for the preparation of nuclear and cytoplasmic extracts as per the manufacturer’s instructions. Cells were washed twice with 1ml of ice-cold PBS/phosphatase inhibitors and 0.3ml of ice-cold PBS/phosphatase inhibitors was added. Cells were removed from the cell culture plate by gentle scraping. Samples were pooled from 2 wells of a 24 well plate. The resulting cell suspension was centrifuged for 5min at 200g in a centrifuge cooled to 4°C. The supernatant was removed, and the pellet kept on ice prior to resuspension in 100ul hypotonic buffer containing 5% (vol/vol) detergent. The resulting cell suspension was centrifuged for 30s at 14,000g in a centrifuge cooled to 4°C. The supernatant was removed as the cytoplasmic fraction. The remaining pellet was resuspended in 25μL of complete lysis buffer and incubated for 30min on ice on a rocking platform. The resulting cell suspension was centrifuged for 10min at 14,000g in a centrifuge cooled to 4°C. The supernatant was removed as the nuclear fraction.

Protein concentration was assessed using a BCA assay kit. 15μg of protein was separated by SDS-page electrophoresis, transferred to a PVDF membrane and blocked in 5% milk/TBS/Tween-20. Immunoblotting was performed by the addition of a primary antibody to protein of interest (antibody details provided in key resource sheet). Proteins were detected with a polyclonal HRP-conjugated secondary antibody and visualized using chemiluminescent detection. Relative protein expression was calculated by normalization to beta actin, beta tubulin or histone 3.

### RT-PCR

RNA was isolated from cells using the RNeasy extraction kit, purity and RNA concentration was assessed using a nanodrop 3000 spectrophotometer. 1μg RNA was reverse transcribed using a SuperScript III First Strand Synthesis system. Resulting cDNA was then diluted 1:20 and amplified using SYBR Green reagents in a StepOne Plus detection system. Cycling conditions: Heat ramp 95°C for 10min, extension (95°C for 15s, 60°C for 1min) for 45 cycles, melt curve 95°C for 15s, 60°C for 1min, 95°C for 15s. Relative gene expression was calculated by normalization to beta actin. The primer sequences used are detailed in key resource sheet.

### Genetic modulation of IL-33

Knock-down of IL-33 from ARPE-19 cells was achieved using the fast-forward transfection technique. Cells were seeded at a concentration of 55,000 per well of a 24 well plate in 0.5ml of culture medium with 1% FCS and no antibiotics. Cells were incubated for 1h at 37°C prior to transfection. The FlexiTube GeneSolution, as a specific mixture of four preselected siRNA duplexes was used to target different sequences of the human Il-33 gene. Each siRNA was diluted in 100μl of culture medium without serum and antibiotics (final concentration 20nM each siRNA). 6μl HiPerfect transfection reagent was added to the siRNA, vortexed and left for 5min. 100μl of transfection complex was added to the cells and left for 48h at 37°C.

For IL-33 overexpression in ARPE-19 cells, a CRISPRcas9 activation plasmid was used to upregulate the expression of the human Il-33 gene. Cells were seeded at a concentration of 40,000 per well of a 24 well plate in 0.5ml of culture medium with 10% FCS and no antibiotics. Cells were incubated overnight at 37°C prior to transfection. Media was replaced just before transfection. For each transfection, 0.16μg of Plasmid DNA was diluted into 25μl plasmid transfection medium. Separately, 0.833μl of transfection reagent was diluted in 25μl plasmid transfection medium. Both solutions were left for 5min before combined, mixed and left for a further 30min. 50ul of transfection complex was added added to the cells and left for 48h at 37°C.

### Cell viability assays

Cell proliferation was assessed using the colorimetric MTT assay as per the manufacturer’s instructions. 10μL of the 12 mM MTT stock solution in 100μL of culture medium was added to cells and incubated at 37°C for 2h. All but 25μL of medium was aspirated from the wells. 50μL of DMSO was added and incubated 37°C for 30min; the solution was transferred into a 96well plate. Absorbances were read at 540nm and expressed as a percentage of untreated control.

Cellular cytotoxicity was quantified using a lactate dehydrogenase (LDH) detection kit as per the manufacturer’s instructions. Briefly, 50μL of cell culture supernatant was transferred to a 96well plate with 50μL of the reaction mixture and incubated 37°C for 30min. Absorbances were read at 450nm and expressed as relative to untreated control.

### Enzyme-based metabolic assays

Glucose consumption was assessed using a glucose coulometric assay kit as per the manufacturer’s instructions. 15μL of cell culture supernatant was transferred to a 96well plate with 85μL of diluted assay buffer. 100μL of enzyme buffer was then added and incubated at 37°C for 10min. Absorbances were read at 520nm. Glucose consumption was calculated by subtracting the glucose concentration of samples from the glucose concentration of unconditioned media and expressed as relative to an untreated control.

Extracellular lactate production was measured using a L-lactate fluorometric assay kit as per the manufacturer’s instructions. 20μL of cell culture supernatant was transferred to a 96well plate with 100μL diluted assay buffer, 20μL of cofactor mixture, 40μL of enzyme mixture and 20μL of fluorometric substrate. The plate was incubated for 20min at RT and read using an excitation wavelength at 530nm and an emission wavelength of 590nm. Lactate production was calculated by subtracting the lactate concentration of samples from the lactate concentration of unconditioned media and expressed as relative to an untreated control.

ATP and ADP measurements were quantified using an ATP/ADP fluorometric assay kit. Cell supernatants were removed and 500μL of nucleotide releasing buffer was added to cells. 90μL of this buffer was transferred to a 96well plate and background ATP fluorescence was measured. 10μL of ATP-monitoring enzyme was added and left for 2min at RT. ATP fluorescence levels were then measured. The fluorescence levels were subsequently measured again to give background ADP fluorescence. 10μL of ADP converting enzyme was then added and left for 2min at RT. ADP fluorescence levels were then measured. ATP and ADP levels were calculated and expressed as the ratio ATP/ADP. Data is expressed as relative to an untreated control.

### ELISA

Cell culture supernatants were analyzed for malonyl CoA using a commercially available ELISA kit (CUSABIO).

### Extracellular flux analysis

Cell metabolism was assessed using a Seahorse XFp Extracellular Flux Analyzer. ARPE-19 and M1-MO were seeded at a density of 30,000 cells per well, 24h prior to analysis, with further treatments detailed in results. Cell-Tak was used to attach primary RPE and BMMC at a density of 50,000 to the XFp. Real time measurements of oxygen consumption rate (OCR) and extracellular acidification rate (ECAR) were normalized to total protein content using a BCA assay. Pre-optimised injections specific for each assay were used. Cell Mito Stress kit injections: oligomycin (1μM), carbonyl cyanide 4-(trifluoromethoxy)phenylhydrazone (FCCP) (0.5μM) and antimycin A/rotenone (1μM). Glycolysis stress injections: glucose (10mM), oligomycin (1μM) and 2-deoxyglucose (2-DG) (1μM). Glycolysis rate injections: antimycin A/rotenone (1μM) and 2-DG (1μM). UK5099 (5μM) was used to inhibit the mitochondrial pyruvate transporter MCP1. Etomoxir (3μM) was used to inhibit CPT1. GSK 2837808A (1μM) was used to inhibit LDHA. Potassium dichloroacetate (25mM) was used to inhibit PDK1.

XF medium included 25mM glucose, 1mM pyruvate and 2mM glutamine in minimal DMEM at pH 7.4. Metabolic parameters were calculated using the following formulae: OCR/ECAR (first OCR measurement)/(first ECAR measurement), proton leak (difference between OCR after oligomycin and non-mitochondrial respiration), non-mitochondrial respiration (OCR measurement after antimycin-A/ rotenone), basal respiration (difference between OCR before oligomycin and non-mitochondrial respiration), ATP production (difference between basal respiration and proton leak), maximal respiration (difference between maximum OCR post FCCP injection and non-mitochondrial respiration), spare respiratory capacity ((maximal respiration) / (basal respiration) x 100), glycolysis (difference between maximum ECAR rate before oligomycin and non-glycolytic acidification), glycolytic capacity (difference between maximum ECAR rate after oligomycin and non-glycolytic acidification), glycolytic reserve (difference between glycolytic capacity and glycolysis), non-glycolytic acidification (last ECAR measurement before glucose injection), basal glycolysis (last glycoPER measurement before rotenone/antimycin-A), compensatory glycolysis (maximum glycoPER measurement after rotenone/antimycin-A), mitoOCR/glycoPER ([(last OCR before rotenone/antimycinA)-(minimum OCR after rotenone/antmycinA)] / basal glycolysis), percentage PER from glycolysis ((basal glycolysis/basal PER) x 100), MPC-dependent respiration (difference between OCR before and after UK5099) and CPT1-dependent respiration (difference between OCR before and after etomoxir).

### Mass spectrometry

Gas chromatography coupled to mass spectrometry (GCMS) was performed using previously detailed methods [54]. For stable isotope tracing (SITA) experiments, ARPE-19 cells were seeded at a concentration of 1,000,000 per well of a 6-well plate in 3ml of normal culture medium with 10% FCS and antibiotics and left for 24h to reach sub-confluency before treatment. Cells were washed twice with PBS before pulsing with with [U-13C]-glucose medium for 2h. SITA media contained 10mM [U-13C]-glucose, 2mM glutamine and 10% dialysed FBS. Cells were washed twice with saline and lysed with 0.8ml methanol, sonicated and freeze-dried using a speed-vacuum. Mass isotopomer distributions (MID) were derived using an algorithm developed at McGill University [55]. Relative metabolite levels were normalised to protein content (μg) using a BCA assay.

### Electron microscopy

For ex vivo EM, WT and Il-33−/− mice eyes were enucleated with the anterior chamber removed and fixed in 2.5% glutaraldehyde in 0.1M phosphate buffer. Eyes were washed in buffer post fixation in Osmium tetroxide for 1 hour and enbloc stained with uranyl acetate prior to being dehydrated with ethanol and embedded in Epon-resin. After polymerisation the embedded eyes were trimmed to remove excess material revealing the retina and choroid layers these were sectioned transversely at 0.5μm for light microscopy. Suitable areas were selected for transmission electron microscopy and these were sectioned (80nm thick) and sectioned stained with uranyl acetate and lead citrate prior to observation in a Tecnai 12 microscope. For in vitro EM, cells were grown to sub confluence in 24 well plates and treated as detailed in the results. Cells were washed in buffer and fixed in phosphate buffered glutaraldehyde and postfixed using osmium tetroxide in the same buffer. Cells were stained with uranyl acetate and then after ethanol dehydration were infiltrated with Epon resin mix (TAAB labs) and polymerised at 60deg C for 2 days. Sections were cut at 70-80nm thickness using an Ultracut S ultramicrotome and stained with Reynolds’ lead citrate and uranyl acetate. Sections were viewed and images recorded on a Tecnai T12 microscope.

### Quantification and Statistical Analysis

#### ImageJ

Mitochondrial morphology was manually assessed using ImageJ software. A scale was set by normalising the pixel distance to the scale provided (in nm) on each image (Figure S16A). Area measurements were calculated by freehand selection around every whole mitochondrion observed in each image (Figure S16B). Three separate measurements were taken the mean value was used as the area. Diameter measurements were calculated by freehand lines at the largest part of every whole mitochondrion observed in each image (Figure S16C). Mitochondrial numbers were manually counted in each image which corresponded to a 4.95μm2 (ex vivo) or 37μm2 (in vitro) area of the RPE; only whole mitochondria were counted (Figure S16D). Mitochondria were manually characterized into either (Figure S16D) !short tubular”, (Figure S16E) !fragmented” or (Figure S16F) !long tubular” phenotypes.

### Seahorse analysis

As the number of wells used of a seahorse plate vary between experiment, the number of technical repeats (wells used) is reported in each experiment. The mean of these technical repeats (wells used) was subsequently used to provide the biological repeat for that experiment. If no technical repeats were used (only one well per condition), that sole value is used for the biological repeat of that experiment. The reported data is of biological repeats from ≤3 independent experiments.

### Statistics

Data are presented as mean ± SD for technical replicates or mean ± SEM for biological replicates. Statistical analysis was performed using an unpaired two-tailed Student’s t test or ANOVA as specified for comparison between groups. P < 0.05 was considered significant.

### Study approval

All procedures were conducted under the regulation of the United Kingdom Home Office Animals (Scientific Procedures) Act 1986 and followed the Association for Research in Vision and Ophthalmology (ARVO) statement for the use of animals in ophthalmic and vision research. The methods were carried out in accordance with the approved University of Bristol institutional guidelines and all experimental protocols under Home Office Project Licence 30/3281 were approved by the University of Bristol Ethical Review Group.

## Acknowledgements

This work was supported by the Macula Society (ADD, ST), the Academy of Medical Sciences (ST), and the National Eye Research Centre (ST, ADD). EEV is funded by an RD Lawrence, Diabetes UK Fellowship (17/0005587).

## Authors contributions

Conceptualization, S.T and A.D.D.; Methodology, S.T. and L.S.; Investigation, L.S., E.V., N.H., J.L., C.N., E.D., and S.T.; Writing – Original Draft, S.T., L.S. and A.D.D.; Writing – Review & Editing, M.C., A.H., E.V., N.H., J.L., C.N., and E.D.; Funding Acquisition, S.T. and A.D.D.; Resources, M.C., E.V., E.D., J.L, and C.N.; Project Management, S.T., Supervision, S.T. and A.D.D.

## Supplemental Information

Supplemental Information includes 16 figures

