## supplementary figures 1-16 for "Interleukin-33 regulates metabolic reprogramming of the retinal pigment epithelium in response to immune stressors"

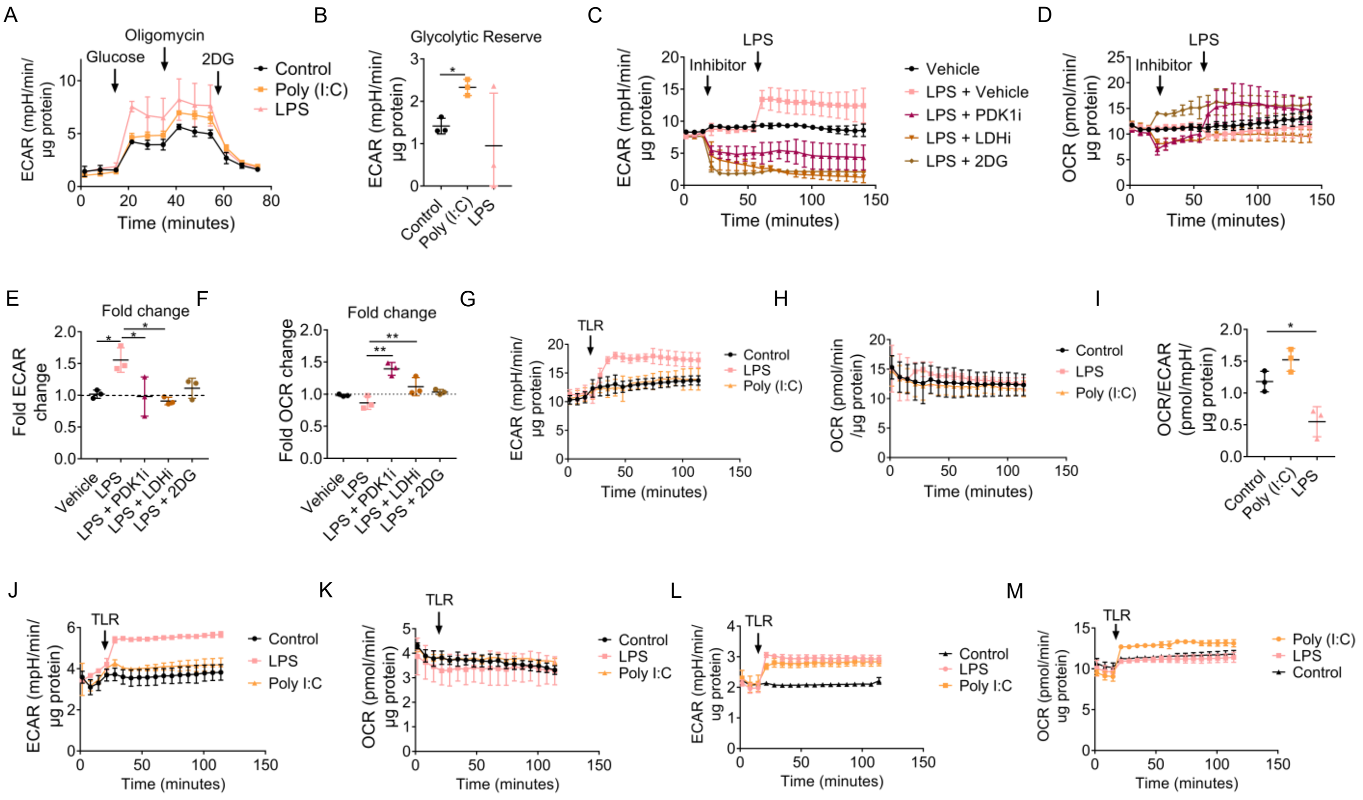


**Figure S1.** (A) Glycolysis stress analysis of ARPE-19 stimulated with either 30min LPS (1µg/ml) or 6h Poly (I:C) (10µg/ml) (*n=3*). (B) Parameters calculated from (A) (as detailed in methods) (*n=3*). Real-time changes in (C) ECAR or (D) OCR following injection of either 2DG (1µM), GSK (LDHAi; 40µM) or DCA (PDKi; 20mM) and following injection of LPS (1µg/ml) to ARPE-19 cells (*n=3*). (E) Fold ECAR change upon LPS stimulation (*n=3*). (F) Fold OCR change upon LPS stimulation (*n=3*). (G) Real-time changes in ECAR following injection of LPS (1µg/ml) or Poly (I:C) (10µg/ml) to primary murine RPE (*n=3*). (H) Real-time changes in OCR following injection of LPS (1µg/ml) or Poly (I:C) (10µg/ml) to primary murine RPE (*n=3*). (I) Basal OCR/ECAR ratio of primary RPE cells stimulated with either 30min LPS (1µg/ml) or 6h Poly (I:C) (10µg/ml) (*n=3*). (J) Real-time changes in ECAR following injection of either LPS (1µg/ml) or Poly (I:C) (10µg/ml) to MIO-M1 cells (*n=2*). (K) Real-time changes in OCR following injection of LPS (1µg/ml) or Poly (I:C) (10µg/ml) to MIO-M1 cells (*n=2*). (L) Real-time changes in ECAR following injection of LPS (1µg/ml) or Poly (I:C) (10µg/ml) to BMMC (*n=2*). (M) Real-time changes in OCR following injection of LPS (1µg/ml) or Poly (I:C) (10µg/ml) to BMMC (*n=3*). Data are expressed as means ± SD from (J-M) two or (A-I) three independent experiments. (A-B) Represents the biological repeats from three independent experiments; each biological repeat is the mean of two technical repeats (two seahorse wells per experiment. (C-F) Represents the biological repeats from three independent experiments; each biological repeat is either the mean of two technical repeats or a single technical repeat (one or two seahorse wells per experiment). (J-M) Represents the biological repeats from two independent experiments; each biological repeat is the mean of two technical repeats (two seahorse wells per experiment. One-way ANOVA with Dunnet's multiple comparisons test; **p*<0.05.

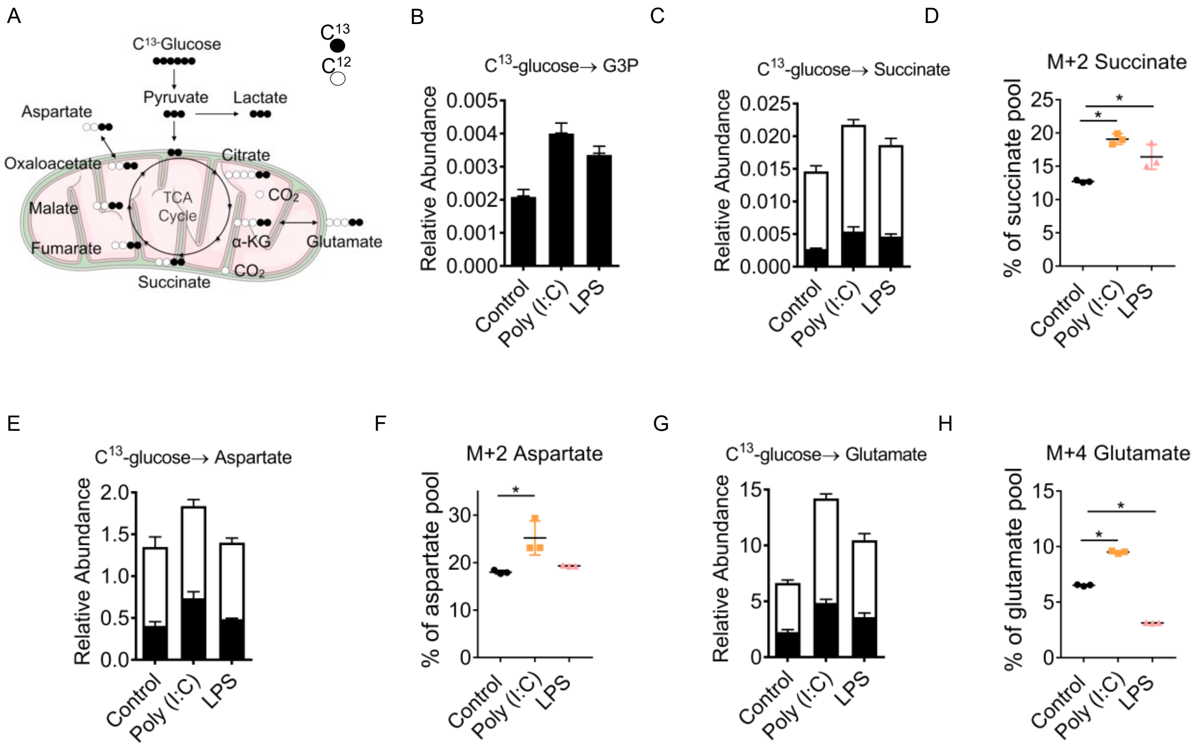


**Figure S2.** (A) Schematic of glucose metabolism in the first turn of the TCA cycle. (B-H) ARPE-19 were treated for either 30min LPS (1μg/ml) or 24h Poly (I:C) (10μg/ml). Uniformly labelled $^{13}$C-glucose incorporation into ARPE-19 metabolites. Relative abundance of $^{12}$C and $^{13}$C including (B) glyceraldehyde-3-phosphate, (C) succinate, (E) aspartate and (G) glutamate. Mass isotopomer distribution of C$^{13}$-glucose-derived carbon into (D) M+2 succinate, (F) M+2 aspartate and (H) M+4 glutamate (*n=3*). (D, F and H) One-way ANOVA with Dunnet's multiple comparisons test; **p*<0.05 Represents data from three independent experiments.

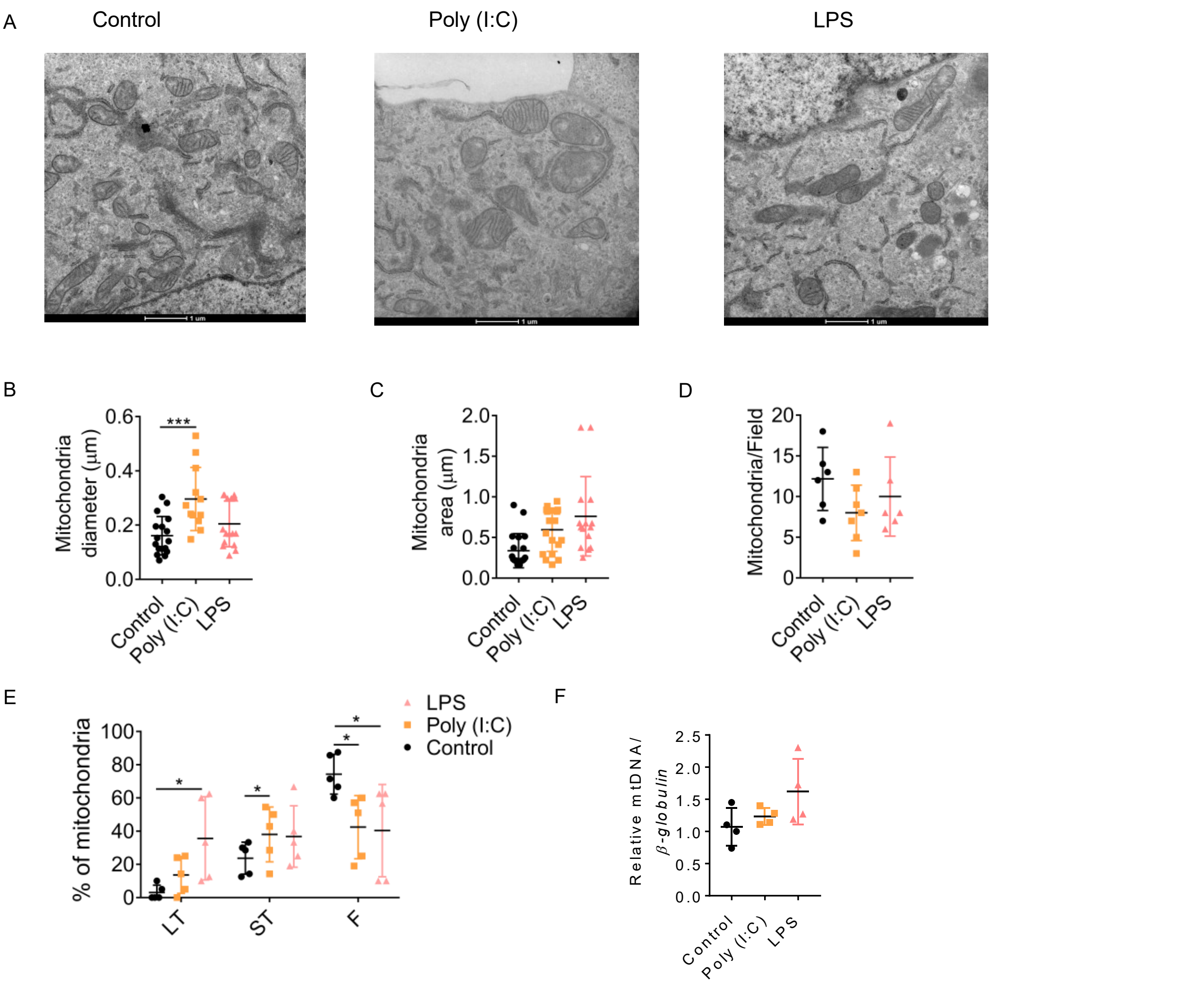


**Figure S3.** (A) Representative transmission electron microscopy of ARPE-19 mitochondria either untreated, stimulated with Poly (I:C) (10µg/ml) or LPS (1µg/ml) for 24h. Magnification 9300x. (B) Mitochondrial diameter, (C) mitochondrial area and (D) mitochondrial number were calculated using ImageJ from transmission electron microscopy images of RPE from ARPE-19 mitochondria either untreated, stimulated with Poly (I:C) (10µg/ml) or LPS (1µg/ml) for 24h. (E) Quantification of mitochondrial morphology into either fragmented, short tubular or long tubular phenotypes. (F) Quantitative RT-PCR was performed to assess mitochondrial content; this was estimated from the amplification of *CTYB* and 16S rRNA relative to *BGLOB* mRNA transcripts. Represents data from (A-E) three and (F) four independent experiments. One-way ANOVA with Dunnet's multiple comparisons test; **p*<0.05, ****p*<0.005.

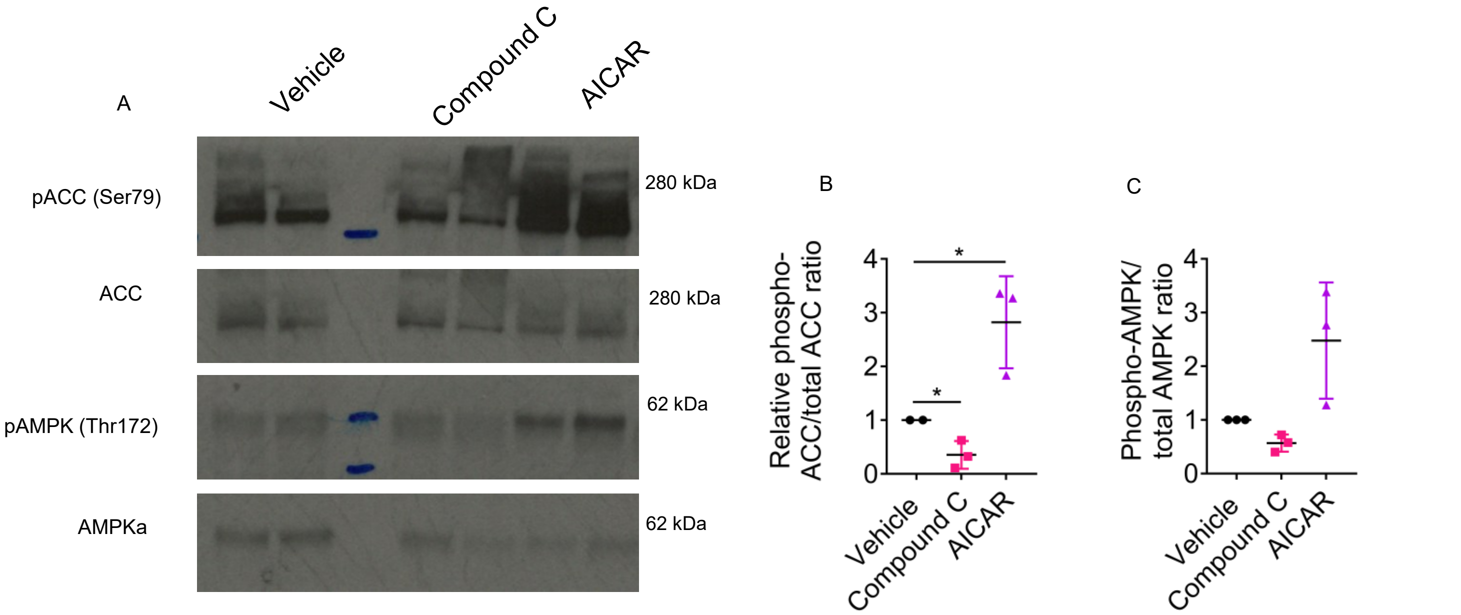


**Figure S4.** (A) Immunoblot of AMPK activation for ARPE-19 treated with either AICAR (1mM) or Compound C (40$\mu$M) for 2h. Represents data from three independent immunoblots. (B) Quantification of phospho-ACC/ACC protein levels (*n=3*). (C) Quantification of phospho-AMPK/AMPK protein levels (*n=3*).

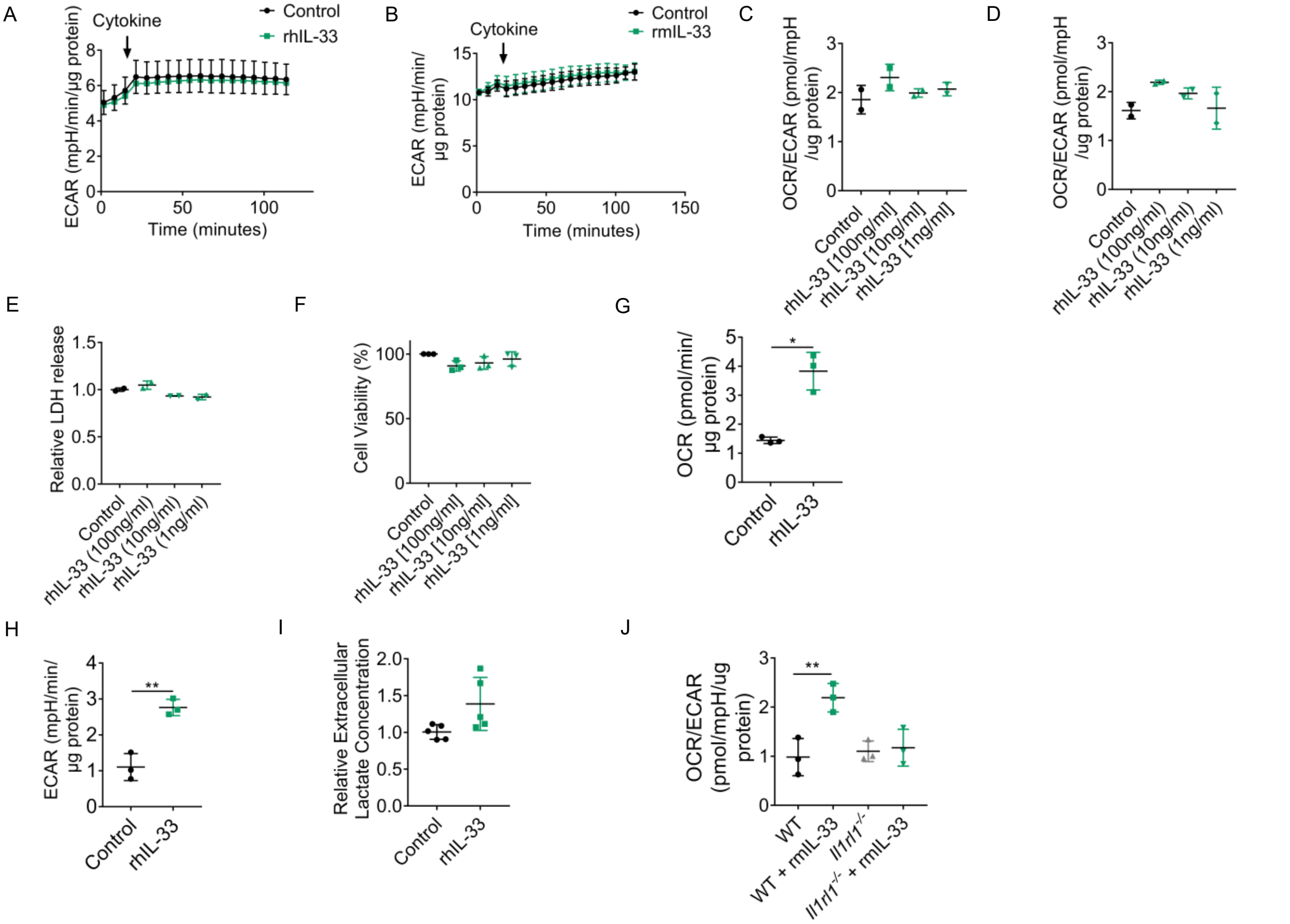


**Figure S5.** (A) Real-time changes in ECAR following injection of rhIL-33 (100ng/ml) to ARPE-19 (*n=3*). (B) Real-time changes in ECAR following injection of rmIL-33 (100ng/ml) to primary murine RPE (*n=3*). (C) Basal OCR and ECAR from ARPE-19 treated with varying doses of rhIL-33 (100, 10 and 1ng/ml) for 12h (*n=3*). (D) Basal OCR and ECAR from ARPE-19 treated with varying doses of rhIL-33 (100, 10 and 1ng/ml) for 24h (*n=2*). (E) LDH release from ARPE-19 treated with varying doses of rhIL-33 (100, 10 and 1ng/ml) for 24h (*n=2*). (F) MTT viability assay of ARPE-19 treated with varying doses of rhIL-33 (100, 10 and 1ng/ml) for 24h (*n=3*). (G) Parameters calculated from (Fig. 3A) (as detailed in methods) (*n=3*). (H) Parameters calculated from (Fig. 3B) (as detailed in methods) (*n=3*). (I) Lactate production in ARPE-19 stimulated with rhIL-33 (100ng/ml) for 24h (*n=5*). (J) Basal OCR/ECAR from WT or *Il1rl1*-/- primary murine RPE treated with rmIL-33 (100ng/ml) for 24h (*n=3*). Data are expressed as means ± SD from (C-E) two, (A-B, F-H and J) three and (I) five experiments. (A-B and G-H) Represents the biological repeats from three independent experiments; each biological repeat is the mean of three technical repeats (three seahorse wells per experiment). (C and D) Represents the biological repeats from two independent experiments; each biological repeat is either the mean of two technical repeats or a single technical repeat (one or two seahorse wells per experiment). (J) Represents the biological repeats from three independent experiments; each biological repeat is either the mean of two technical repeats or a single technical repeat (one or two seahorse wells per experiment). (C, D, E, F and J) One-way ANOVA with Dunnet's multiple comparisons test; **$p<0.01$. Unpaired (G, H and I) Student's T-test; *$p<0.05$, **$p<0.01$.

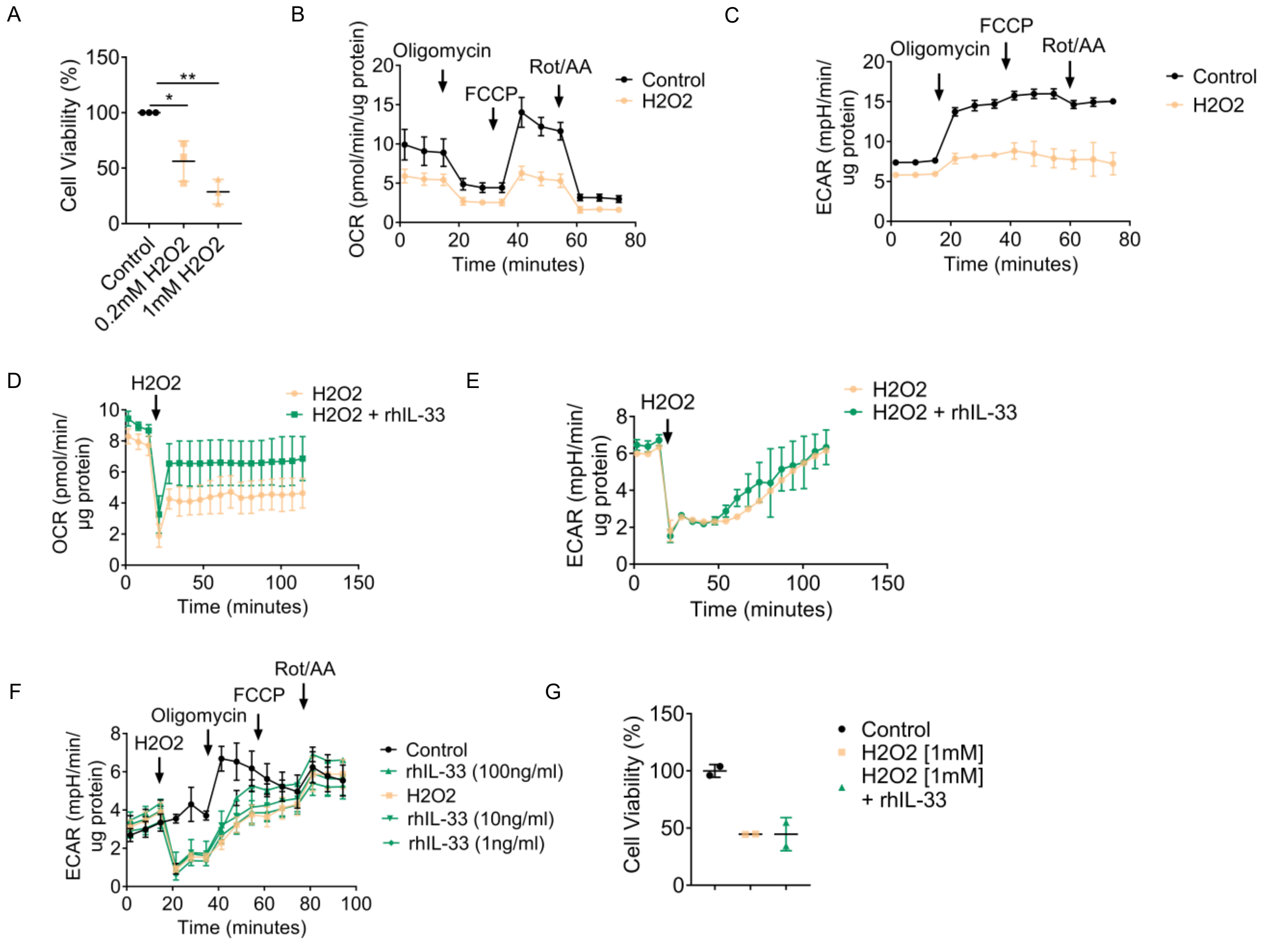


**Figure S6.** (A) MTT viability assay of ARPE-19 treated with varying doses of $H_2O_2$ (1 and 0.2mM) for 24h (*n=3*). (B) A mitochondrial stress test was used to assess mitochondrial function after treatment with $H_2O_2$ (1mM) for 24h (*n=2*). (C) Corresponding ECAR data from (B) (*n=2*). ARPE-19 were pre-treated with rhIL-33 (100ng/ml) for 12h; OCR (D) or ECAR (E) measurements were taken following the injection of $H_2O_2$ (1mM) (*n=3*). (F) Corresponding ECAR data from Figure 3J (*n=3*). (G) MTT viability assay of ARPE-19 treated with either $H_2O_2$ (1mM) or $H_2O_2$ (1mM) and IL-33 (100ng/ml) for 24h (*n=2*). Data are representative of (A,D-E and F) three or (B-C and G) two independent experiments. (A, D and E) Represents the biological repeats from three independent experiments; each biological repeat is the mean of three technical repeats (three seahorse wells per experiment). (B and C) Represents the biological repeats from two independent experiments; each biological repeat is the mean of three technical repeats (three seahorse wells per experiment). (F) Represents the biological repeats from three independent experiments; each biological repeat is either the mean of two technical repeats or a single technical repeat (one or two seahorse wells per experiment).One-way ANOVA with Dunnet's multiple comparisons test; *$p$<0.05, **$p$<0.01.

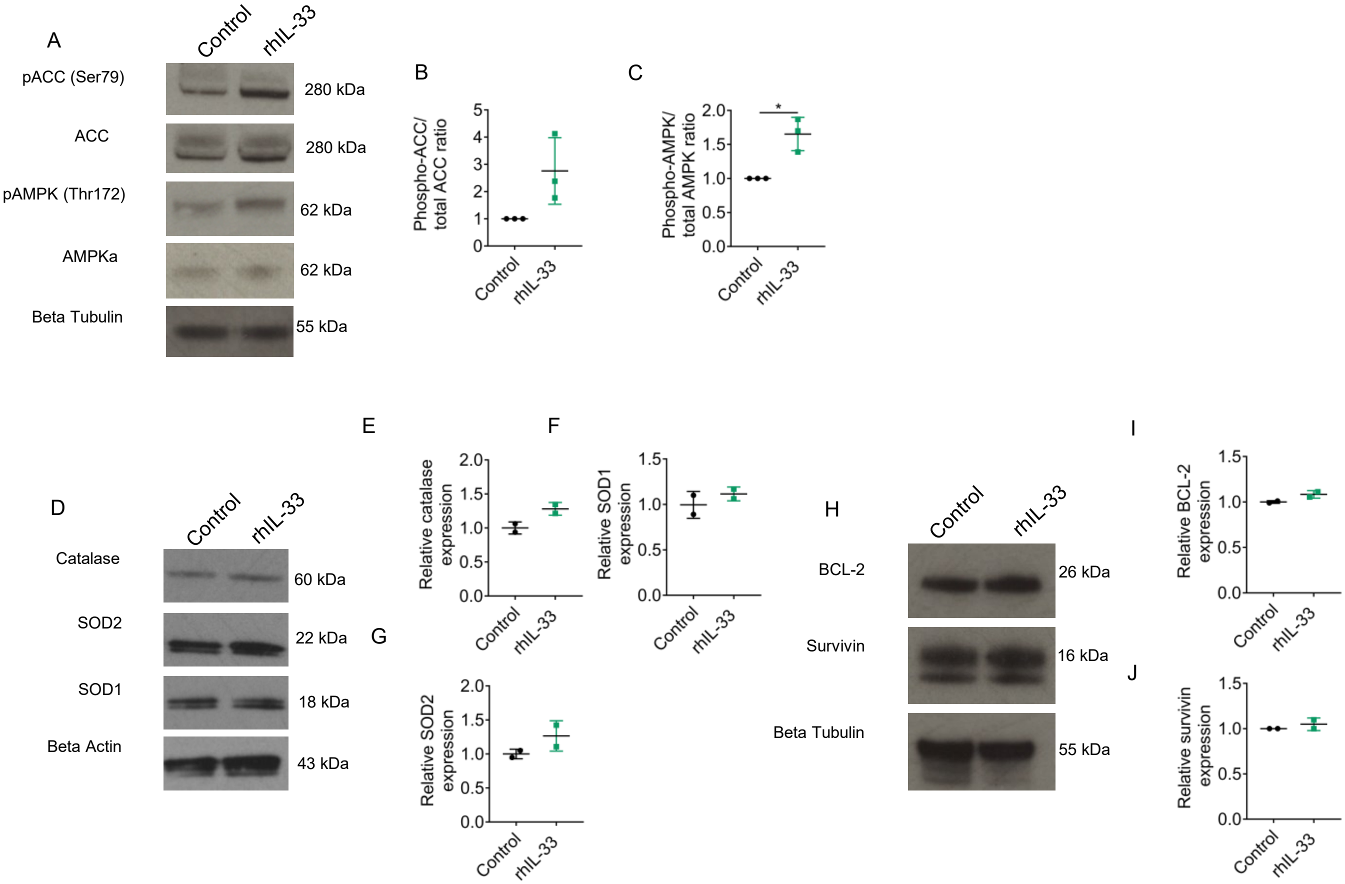


**Figure S7.** ARPE-19 were treated with rhIL-33 (100ng/ml) for 24h; protein was extracted and immunoblot analysis was used to determine: (A-C) the phosphorylation of AMPK and ACC (*n=3*), (D-G) the expression of catalase, SOD1/2 (*n=2*), (H-J) the expression of BCL-2 and surviving (*n=2*). Represents data from three independent blots. (D-G and H-J) Represents data from two independent immunoblots. Unpaired Student's T-test; $*p<0.05$, $**p<0.01$.

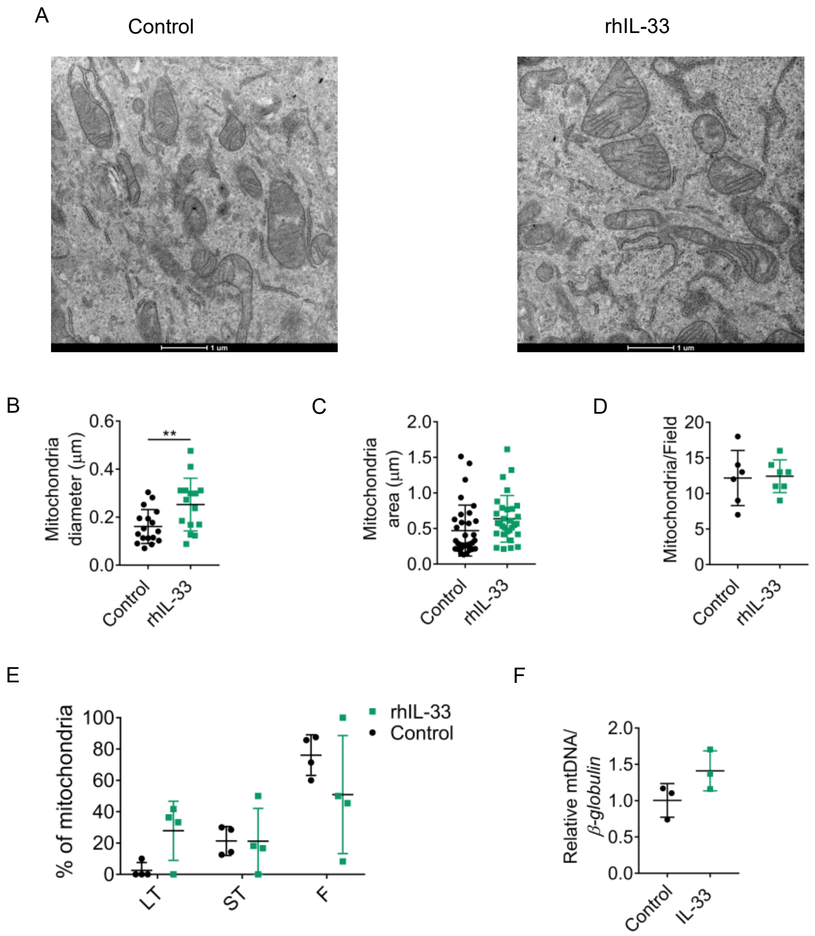


**Figure S8.** (A) Representative transmission electron microscopy of ARPE-19 mitochondria either untreated or stimulated with rhIL-33 (100ng/ml) for 24h. Magnification 9300x. (B) Mitochondrial diameter, (C) mitochondrial area and (D) mitochondrial number were calculated using ImageJ from transmission electron microscopy images of RPE from ARPE-19 mitochondria either untreated or stimulated with rhIL-33 (100ng/ml) for 24h. (E) Quantification of mitochondrial morphology into either fragmented, short tubular or long tubular phenotypes. (F) Quantitative RT-PCR was performed to assess mitochondrial content; this was estimated from the amplification of *CTYB* and *16S rRNA* relative to *BGLOB* mRNA transcripts. Data are representative of three independent experiments. Unpaired Student's T-test; ***p*<0.01.

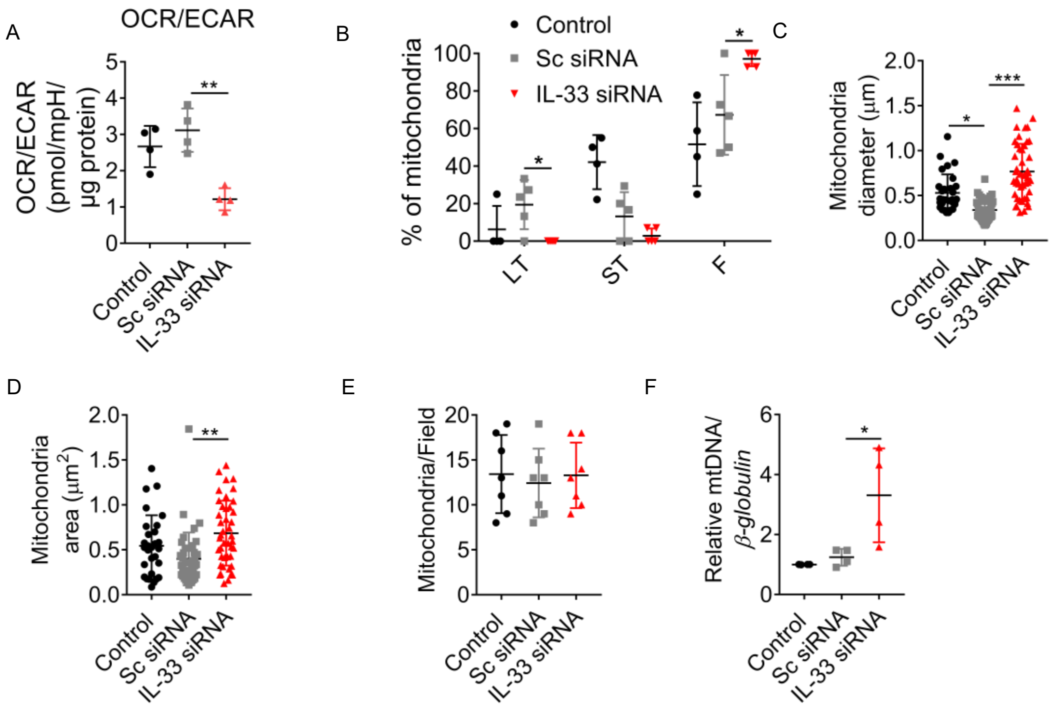


**Figure S9**. (A) OCR/ECAR ratio from ARPE-19 cells transfected with either an IL-33 siRNA or scrambled siRNA (*n=4*). (B) Quantification of mitochondrial morphology into either fragmented, short tubular or long tubular phenotypes (represents data presented in Figure 4L) from ARPE-19 cells transfected with either an IL-33 siRNA or scrambled siRNA. (C) Mitochondrial diameter, (D) mitochondrial area and (E) mitochondrial number were calculated using ImageJ from transmission electron microscopy images of ARPE-19 cells transfected with either an IL-33 siRNA or scrambled siRNA. (F) Quantitative RT-PCR was performed to assess mitochondrial content; this was estimated from the amplification of *CTYB* and 16S rRNA relative to *BGLOB* mRNA transcripts. Data are representative of (A and F) four and (B-E) three independent experiments. (A) Represents the biological repeats from four independent experiments; each biological repeat is the mean of two technical repeats (two seahorse wells per experiment). Unpaired Student's T-test; **p*<0.05, ***p*<0.01. One-way ANOVA with Dunnet's multiple comparisons test; **p*<0.05, ***p*<0.01.

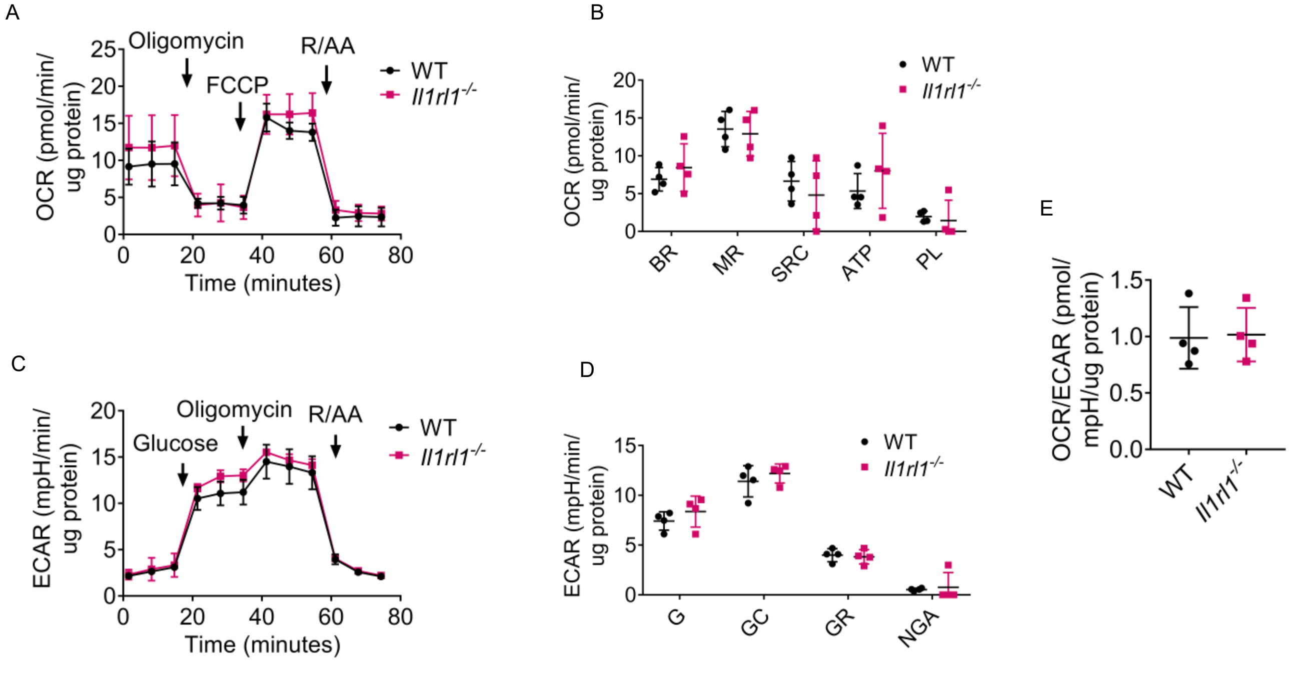


**Figure S10**. Primary RPE were isolated from WT or *Il1rl1-/-* mice. (A) Mitochondrial stress test (*n=4*). (B) Mitochondrial parameters calculated from (A) (*n=4*). (C) Glycolysis stress test (*n=4*). (D) Glycolysis stress parameters calculated from (C) (*n=4*). (E) Basal OCR and ECAR measurements expressed as the ratio OCR/ECAR (*n=4*). Data presented as means ± SD. Represents the biological repeats from three independent experiments; each biological repeat is either the mean of two technical repeats or a single technical repeat (one or two seahorse wells per experiment).

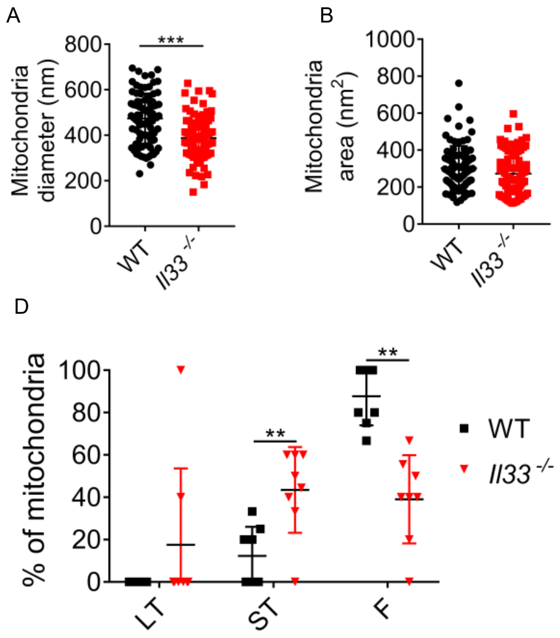


**Figure S11**. Primary RPE were isolated from WT or *Il33-/-* mice. (A) Mitochondrial diameter, (B) mitochondrial area and (C) mitochondrial number were calculated using ImageJ from transmission electron microscopy images of RPE from WT and *Il-33-/-*mice (as detailed in methods). (D) Quantification of mitochondrial morphology into either fragmented, short tubular or long tubular phenotypes. Represents data presented in 5H. Data are expressed as means ± SD from three independent experiments. Unpaired Student's T-test; *$p$<0.05, *$p$<0.01, ***$p$<0.001.

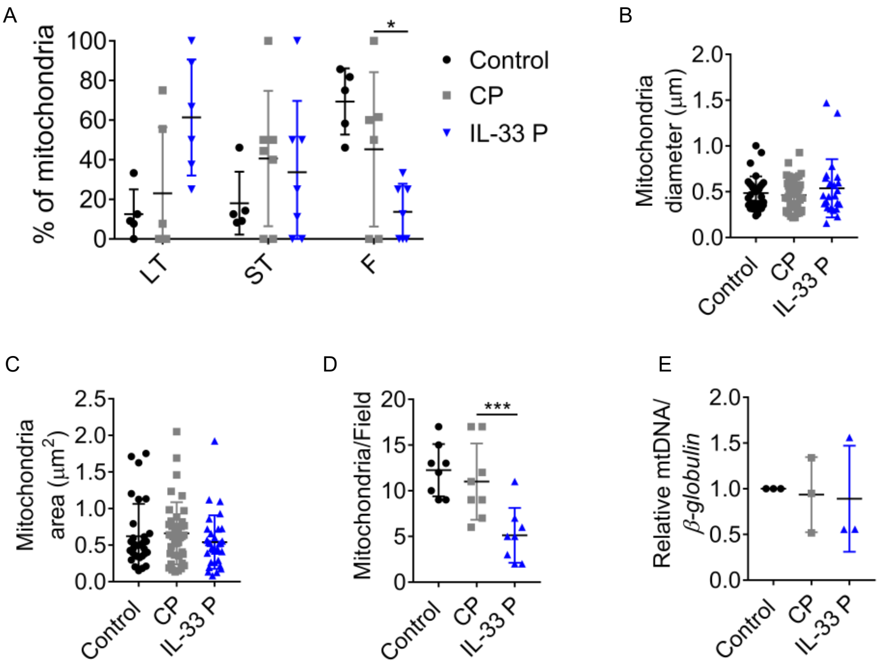


**Figure S12**. ARPE-19 cells transfected with either an IL-33 activation plasmid or scrambled gRNA activation plasmid. (A) Mitochondrial diameter, (B) mitochondrial area and (C) mitochondrial number were calculated using ImageJ from transmission electron microscopy images of RPE (as detailed in methods). (D) Quantification of mitochondrial morphology into either fragmented, short tubular or long tubular phenotypes. Represents data presented in Figure 6. Data are expressed as means ± SD from three independent experiments. One-way ANOVA with Dunnet's multiple comparisons test; *$p$<0.05, ***$p$<0.001.

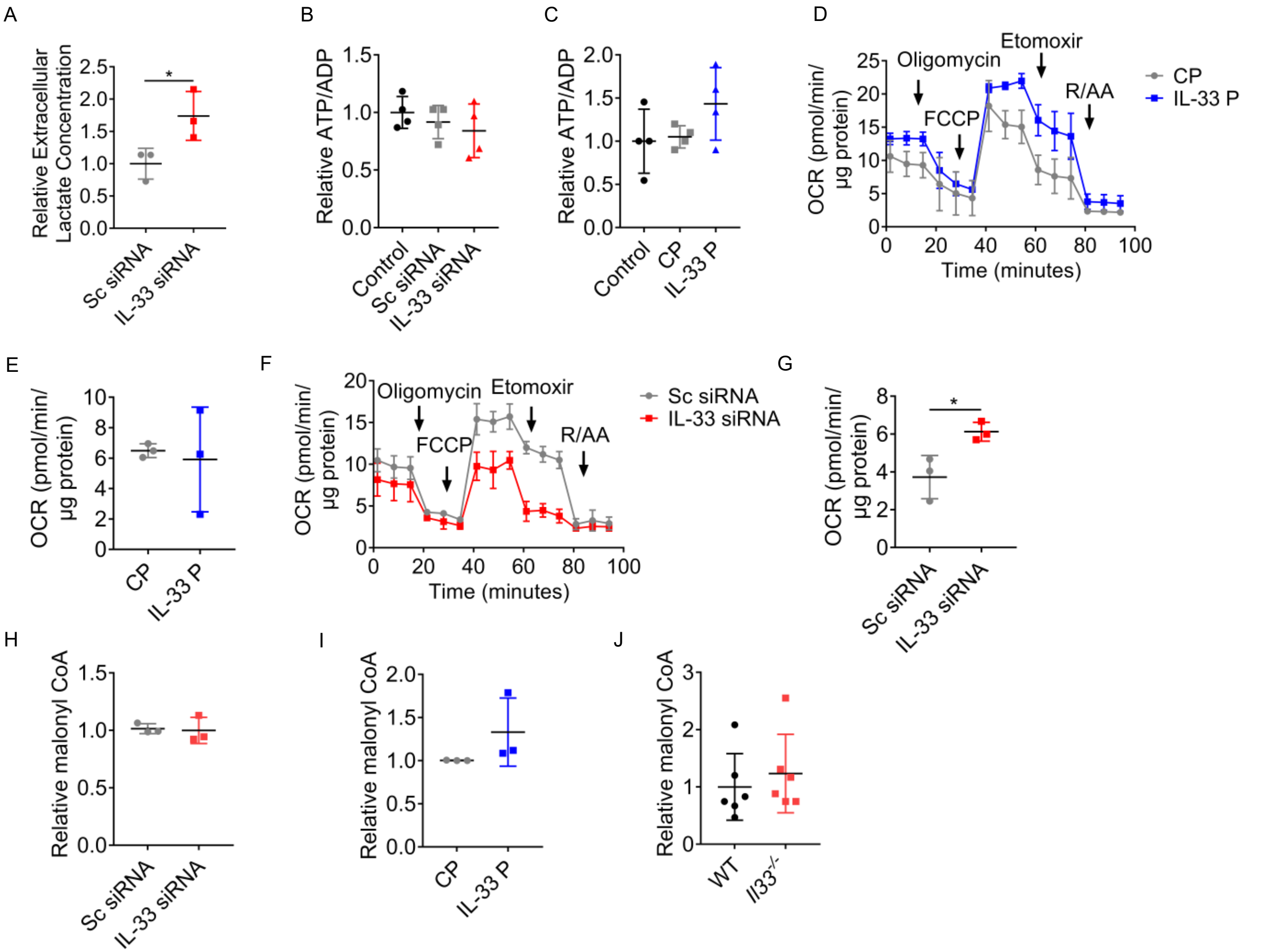


**Figure S13**. (A) Relative lactate extracellular concentration of ARPE-19 transfected with either an IL-33 siRNA or scrambled siRNA (*n=3*). (B) Relative ATP/ADP ratio in ARPE-19 cells transfected with either an IL-33 siRNA or scrambled siRNA (*n=4*). (C) Relative ATP/ADP ratio in ARPE-19 cells transfected with either an IL-33 activation plasmid or a scrambled gRNA activation plasmid (*n=4*). (D) Modified mitochondrial stress test including a third injection of etomoxir (3µg/ml) of ARPE-19 cells transfected with either an IL-33 activation plasmid or a scrambled gRNA activation plasmid (*n=3*). (E) Parameters calculated from (D) (as detailed in methods) (*n=3*). (F) Modified mitochondrial stress test including a third injection of etomoxir (3µg/ml) of ARPE-19 cells transfected with either an IL-33 siRNA or scrambled siRNA (*n=3*). (G) Parameters calculated from (F) (as detailed in methods) (*n=3*). (H) Relative malonyl CoA concentration of ARPE-19 transfected with either an IL-33 siRNA or scrambled siRNA (*n=3*). (I) Relative malonyl CoA in ARPE-19 lysates transfected with either an IL-33 activation plasmid or a scrambled gRNA activation plasmid (*n=3*). (J) Relative malonyl coA levels in RPE lysates from WT and *Il-33-/-* mice (*n=6*). Represents data from (A and D-H) three, (B-C) four and (J) six independent experiments. (D-G) Represents the biological repeats from three independent experiments; each biological repeat is the mean of three technical repeats (three seahorse wells per experiment). Unpaired Student's T-test; *$p$<0.05, **$p$<0.01.

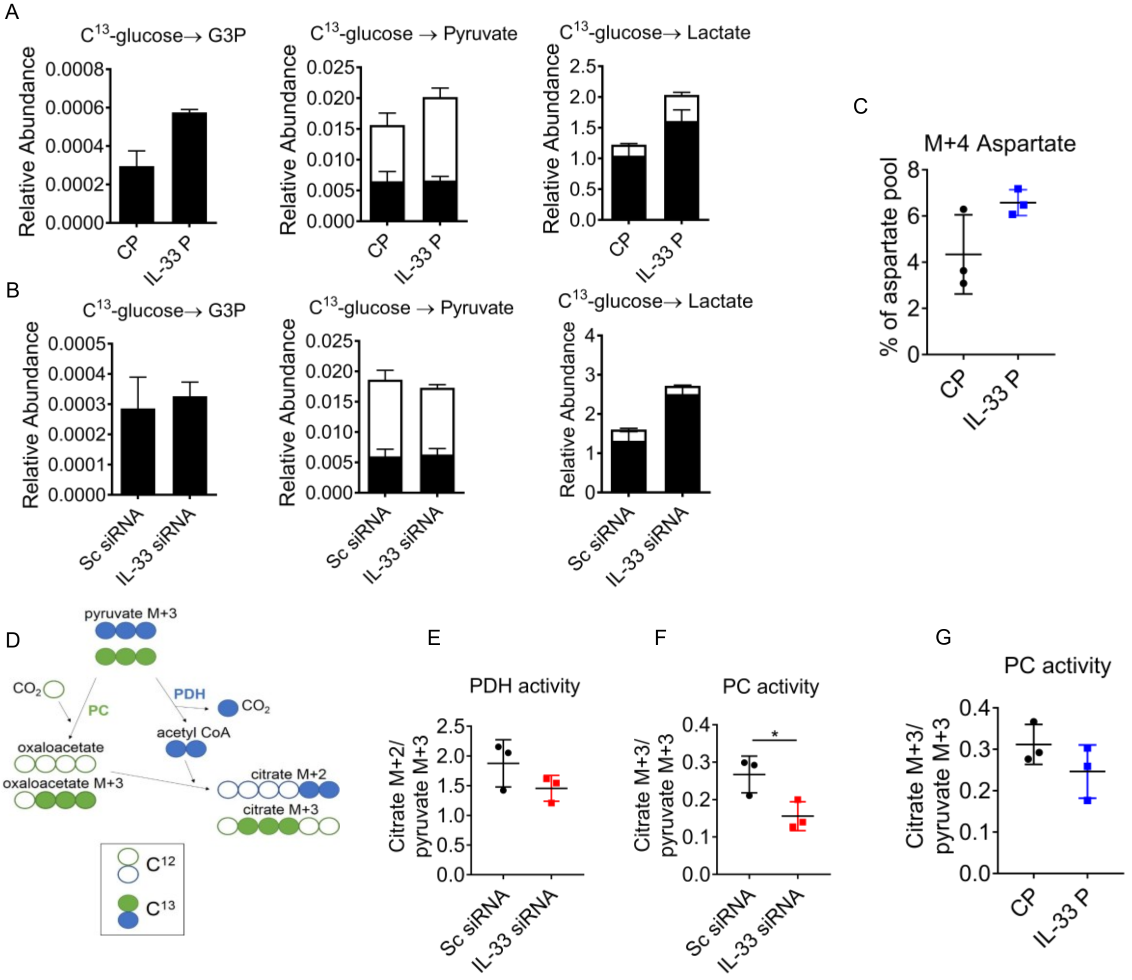


**Figure S14**. (A) ARPE-19 cells were transfected with either an IL-33 activation plasmid or a scrambled gRNA activation plasmid. Uniformly labelled $^{13}$C-glucose incorporation into ARPE-19 glycolytic metabolites. Relative abundance of $^{12}$C and $^{13}$C, including glyceraldehyde-3-phosphate, pyruvate and lactate (*n=3*). (B) ARPE-19 cells were transfected with either an IL-33 siRNA or a scrambled siRNA. Uniformly labelled $^{13}$C-glucose incorporation into ARPE-19 glycolytic metabolites. Relative abundance of $^{12}$C and $^{13}$C including glyceraldehyde-3-phosphate, pyruvate and lactate (*n=3*). (C) Mass isotopomer distribution of C$^{13}$-glucose-derived carbon into aspartate pools (*n=3*). (D) Isotopologue distribution in citrate used to obtain a clearer representation of pathways feeding into the TCA cycle. The citrate M+2/pyruvate M+3 ratio can serve as a surrogate for PDH activity, whereas the citrate M+3/pyruvate M+3 ratio can be a surrogate for PC. (E) Citrate M+2/pyruvate M+3 ratio and (F) citrate M+3/pyruvate M+3 ratio in ARPE-19 cells transfected with either an IL-33 siRNA or a scrambled siRNA (*n=3*). (G) citrate M+3/pyruvate M+3 ratio in ARPE-19 cells transfected with either an IL-33 activation plasmid or a scrambled gRNA activation plasmid (*n=3*). Data are expressed as means ± SD from three independent experiments. Unpaired Student's T-test; *$p<0.05$.

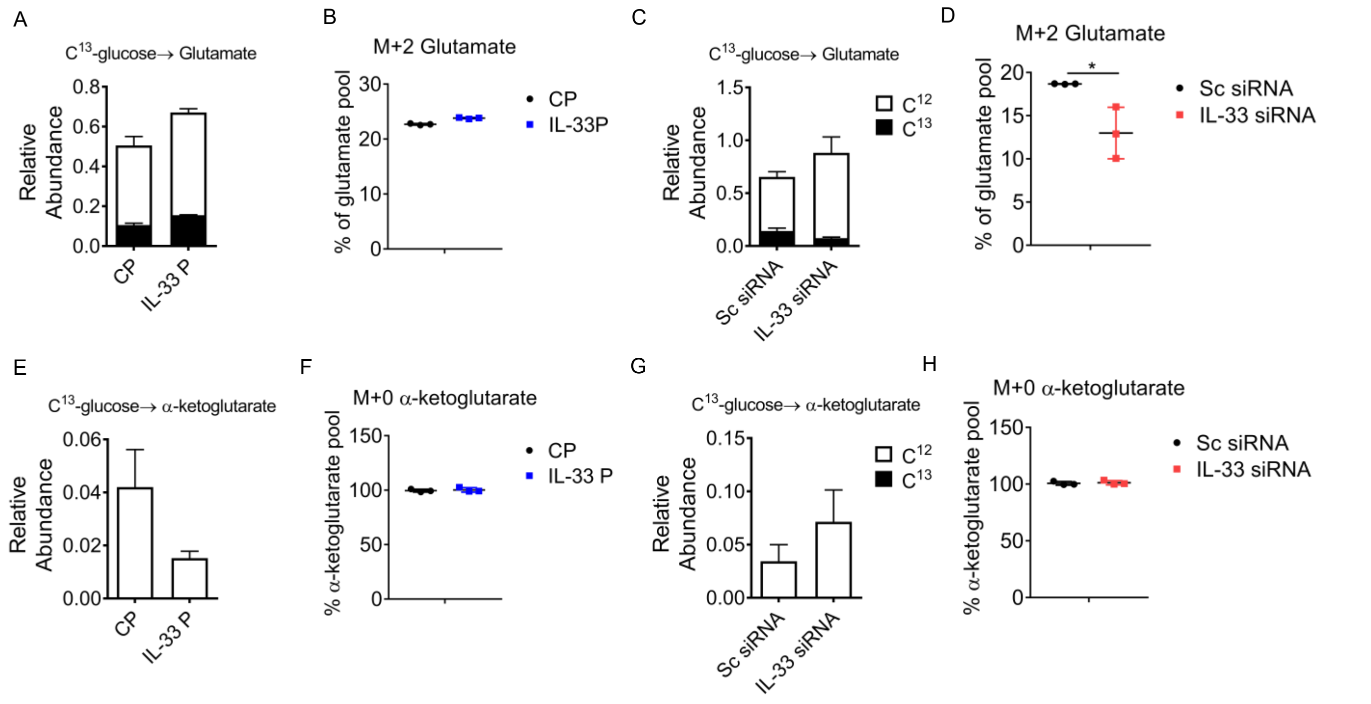


**Figure S15**. (A) ARPE-19 cells were transfected with either an IL-33 activation plasmid or scrambled gRNA activation. Uniformly labelled $^{13}$C-glucose incorporation into glutamate (*n=3*). (B) Mass isotopomer distribution of $C^{13}$-glucose-derived carbon into M+2 glutamate pools (*n=3*). (C) ARPE-19 cells were transfected with either scrambled siRNA or IL-33 siRNA. Uniformly labelled $^{13}$C-glucose incorporation into glutamate (*n=3*). (D) Mass isotopomer distribution of $C^{13}$-glucose-derived carbon into M+2 glutamate pools (*n=3*). (E) ARPE-19 cells were transfected with either an IL-33 activation plasmid or scrambled gRNA activation. Uniformly labelled $^{13}$C-glucose incorporation into α-ketoglutarate (*n=3*). (F) Mass isotopomer distribution of $C^{13}$-glucose-derived carbon into α-ketoglutarate pools (*n=3*). (G) ARPE-19 cells were transfected with either scrambled siRNA or IL-33 siRNA. Uniformly labelled $^{13}$C-glucose incorporation into α-ketoglutarate (*n=3*). (H) Mass isotopomer distribution of $C^{13}$-glucose-derived carbon into α-ketoglutarate pools (*n=3*). Data are expressed as means ± SD from three independent experiments. Unpaired Student's T-test; *$p<0.05$.

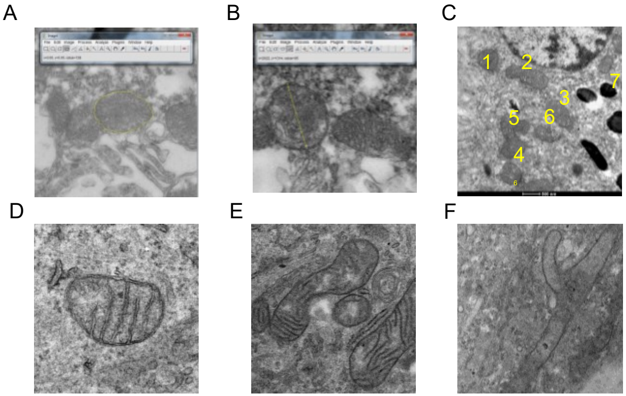


**Figure S16.** (A) Area measurements were calculated by freehand selection around every whole mitochondrion observed. (B) Diameter measurements were calculated by freehand lines at the largest part of every whole mitochondrion observed in each image. (C) Mitochondrial numbers were manually counted in each image. Mitochondria were manually classed into either (D) “short tubular”, (E) “fragmented” or (F) “long tubular“. As detailed in methods.
